# A Foxf1-Wnt-Nr2f1 cascade promotes atrial cardiomyocyte differentiation in zebrafish

**DOI:** 10.1101/2024.03.13.584759

**Authors:** Ugo Coppola, Jennifer Kenney, Joshua S. Waxman

## Abstract

Nr2f transcription factors (TFs) are conserved regulators of vertebrate atrial cardiomyocyte (AC) differentiation. However, little is known about the mechanisms directing Nr2f expression in ACs. Here, we identified a conserved enhancer 3’ to the *nr2f1a* locus, which we call *3’reg1-nr2f1a* (*3’reg1*), that can promote Nr2f1a expression in ACs. Sequence analysis of the enhancer identified putative Lef/Tcf and Foxf TF binding sites. Mutation of the Lef/Tcf sites within the *3’reg1* reporter, knockdown of Tcf7l1a, and manipulation of canonical Wnt signaling support that Tcf7l1a is derepressed via Wnt signaling to activate the transgenic enhancer and promote AC differentiation. Similarly, mutation of the Foxf binding sites in the *3’reg1* reporter, coupled with gain- and loss-of-function analysis supported that Foxf1 promotes expression of the enhancer and AC differentiation. Functionally, we find that Wnt signaling acts downstream of Foxf1 to promote expression of the *3’reg1* reporter within ACs and, importantly, both Foxf1 and Wnt signaling require Nr2f1a to promote a surplus of differentiated ACs. CRISPR-mediated deletion of the endogenous *3’reg1* abrogates the ability of Foxf1 and Wnt signaling to produce surplus ACs in zebrafish embryos. Together, our data support that downstream members of a conserved regulatory network involving Wnt signaling and Foxf1 function on a *nr2f1a* enhancer to promote AC differentiation in the zebrafish heart.

**Author Summary:** Vertebrate hearts are comprised of atrial chambers, which receive blood, and ventricular chambers which expel blood, whose functions need to be coordinated for proper blood circulation. During development, different genetic programs direct the development of these chambers within the vertebrate heart. Members of a family of genes called *Nr2fs* are conserved regulators of atrial chamber development in vertebrates, with mutations in *Nr2f2* of humans being associated with congenital heart defects affecting the atrium. Here, we examine how the gene *nr2f1a*, which is required for normal atrial chamber development in the model zebrafish, is regulated. Using tools, including transgenic reporter lines and genetic mutants, we identify that factors previously shown to regulate atrial chamber development in mammals have conserved roles regulating a genetic element that promotes *nr2f1a* expression within developing atrial cells. Since there is a lack of understanding regarding regulation of *Nr2f* genes during vertebrate atrial cell development, our work provides insights into the conservation of genetic networks that promote heart development in vertebrates and if perturbed could underlie congenital heart defects.

## Introduction

Integrated regulatory networks drive chamber-specific atrial and ventricular cardiomyocyte programs from spatially and temporally defined progenitor populations [1–3]. Key factors within these regulatory networks are nuclear receptor subfamily 2 group F members (Nr2fs; formerly called Coup-tfs), which are critical regulators of vertebrate atrial cardiomyocyte (AC) differentiation during earlier stages of cardiogenesis [4] and subsequently maintain AC identity at later stages [4–6]. Although both Nr2f1 and Nr2f2 paralogs in humans and mice are expressed in ACs [5–7], genetic mapping and functional analysis in cell culture and animal models support that Nr2f2 is the predominant Nr2f that is required for AC differentiation in mammals. In humans, genetic mapping studies have associated mutations in *NR2F2* with structural congenital heart defects (CHDs), including atrioventricular septal defects (AVSDs) [7–9], a malformation predominantly caused by failure of ACs at the venous pole to differentiate and form the atrial septum [10]. Consistent with the necessity of Nr2f2 in mammalian AC differentiation, global *Nr2f2* knockout (KO) mice have small, dysmorphic atrial chambers that lack septa [4]. Furthermore, NR2F2 has a greater requirement promoting atrial differentiation compared to NR2F1 in human embryonic and induced pluripotent stem cell-derived models [11]. In contrast to mammalian Nr2fs, zebrafish *nr2f1a* is the functional homolog of mammalian Nr2f2, as it is required for AC differentiation and the maintenance of AC identity [12,13]. Hence, loss of *nr2f1a* results in zebrafish embryos with small atria reminiscent of the atrial defects of Nr2f2 global KO mice [4,11].

While studies in human stem cell-derived cardiomyocytes and mice cumulatively have demonstrated that Nr2f2 is a critical node within a regulatory network that simultaneously promotes atrial differentiation and represses ventricular differentiation within cardiomyocytes [4,5,11,14], we have limited understanding of the upstream regulation of *Nr2f* genes that lead to the proper differentiation of vertebrate ACs, with a variety of somewhat disparate signals and factors showing they can regulate Nr2f expression. In many *in vivo* and *in vitro* developmental contexts, retinoic acid (RA) signaling, which is required to limit heart size and promote differentiation of the posterior second heart field (SHF) [15–18], is necessary and sufficient to promote *Nr2f1* and *Nr2f2* expression [19–22]. Furthermore, RA signaling is sufficient to induce AC differentiation in many *in vitro* induced pluripotent stem cell protocols [23–25]. However, despite candidate RA response elements and epigenetic changes near the promoter [22], it has not been shown if RA signaling directly regulates *Nr2f* expression within differentiating ACs [26]. In human embryonic stem cell-derived cardiomyocytes, the TF Isl1, which is required for SHF differentiation, negatively regulates *NR2F1* downstream of RA signaling [27]. Additionally, Tbx20, a conserved regulator of cardiomyocyte proliferation [28], is able to promote *Nr2f2* in the ACs of mice through an evolutionarily conserved enhancer [28]. However, as *Isl1* and *Tbx20* are expressed more broadly, and not specifically expressed in differentiating AC progenitors at the venous pole or differentiated ACs, how these TFs coordinate with other signals to promote Nr2f2 expression in ACs is not clear. Similarly, Hedgehog (Hh) signaling has been shown to be able to promote mouse *Nr2f2* via Gli TFs binding to its promoter *in vitro* [29], although this has not been shown in the heart development. Taken together, it is interesting that many of these factors that regulate Nr2fs, in particular RA and Hh signaling, participate in a regulatory network that controls the timing of SHF progenitor differentiation at the venous pole [30,31]. Elegant work from murine and stem cell models has demonstrated that Hh signaling is an upstream factor in a regulatory network involving RA and Wnt signaling, Foxf1/2 and Tbx5 TFs that controls the proper timing of SHF progenitor differentiation at the venous pole of the heart [30,32]. Loss of most of these factors results in AVSDs in murine models and humans [33,34], again consistent with the failure of proper AC differentiation and the impairment of dorsal mesenchymal protrusion development, which contributes to the atrial septum. Despite this understanding of upstream factors in this regulatory network directing AC differentiation at the venous pole in mammals, the mechanisms promoting Nr2f expression in ACs and if there is a connection to the Hh-dependent regulatory network controlling posterior SHF addition are not understood.

Here, we sought to better understand regulatory networks directing Nr2f expression in the differentiation of ACs. Examining regions of open chromatin from Assay for Transposase-Accessible Chromatin (ATAC)-seq analysis of embryonic ACs [35] that were conserved within vertebrates [13], we identified an enhancer 3’ to the *nr2f1a* locus, which we named *3’reg1-nr2f1a* (*3’reg1*), that was able to promote expression within ACs of stable transgenic lines. A Lef/Tcf binding site and multiple Foxf binding sites were found within the enhancer. Mutation or deletion of the Left/Tcf site led to ectopic *3’reg1* reporter activation within the heart, while mutation or deletion of the Foxf sites led to decreased *3’reg1* reporter expression within ACs. Complementary gain- and loss-of-function experiments manipulating Wnt signaling and Foxf1 coupled with epistasis analysis support a model whereby Wnt signaling, which functions at the level or downstream of Foxf1, relieves Tcf7l1a-mediated repression of the *3’reg1* enhancer and promote AC differentiation. However, in contrast to the established role of Hh signaling functioning upstream of these factors in venous SHF differentiation in mammals, Hh was found to modestly repress transgenic *3’reg1* reporter expression and did not affect AC differentiation. Consistent with Nr2f1a functioning downstream of Wnt signaling and Foxf1, neither is sufficient to promote a surplus of ACs in zebrafish *nr2f1a* mutants. Furthermore, CRISPR-mediated deletion of the endogenous *3’reg1* enhancer showed that this abrogated the ability of Wnt signaling and Foxf1 to enhance the number of differentiated ACs. Cumulatively, our results support that downstream components of a regulatory network involving Wnt signaling and Foxf1, which control differentiation of the SHF in mammals, promote AC differentiation via directing zebrafish *nr2f1a* expression through a conserved 3’ enhancer.

## Results

### A *nr2f1a* enhancer drives expression in ACs

To decipher regulatory networks that control *Nr2f* gene expression with respect to its role in heart development, we first examined ∼1.5 kb upstream of the transcription start site (TSSs) of *nr2f1a* as well as the 5’-untranslated region (5’-UTR) for the ability to promote expression in ACs (**S1A Fig. and S1 Table**). Only 1.5 kb upstream were chosen as to not include an adjacent lincRNA, which could complicate analysis. The 5’-UTR was included because we previously identified a conserved RA response element in the *nr2f1a* 5’-UTR [13]. Five different constructs encompassing the 1.5 kb of the *nr2f1a* promoter and different lengths of 5’-UTR were examined with reporter vectors driving GFP (**S1A-F Fig.**). In stable transgenic lines, all the cloned regions showed similar broad expression throughout the embryos, including in the nervous system, somites, and the heart (**S1B-F Fig.**). Additionally, the promoter constructs were not responsive to manipulation of RA signaling (data not shown). Thus, regulatory elements contained within the proximal promoter do not appear to drive *nr2f1a* expression specifically in ACs, consistent with the broader transcriptional regulation generally found with proximal promoters [36], and more distal *cis*-regulatory elements (CREs) likely promote AC-specific *nr2f1a* expression in zebrafish.

Given the conservation of their expression within ACs, we next postulated that *Nr2f* genes might have conserved CREs that promote their AC expression. To identify conserved CREs that promote expression specifically in the ACs, we examined regions of open chromatin found in our recently reported ATAC-seq on isolated ACs from 48 hours post-fertilization (hpf) embryos [35] for conservation using genomic alignments of all vertebrate *Nr2f1* loci with VISTA plots (**Fig. 1A-C and S2A,B,D,F,H Fig.**). Although there is a conserved gene desert 5’ to vertebrate *Nr2f* genes, we did not find conservation of sequences within regions of open chromatin in the ∼700kb 5’ distal the zebrafish *nr2f1a*. The regions of open chromatin with conserved sequence in vertebrate genomes were primarily within ∼20kb 5’ and 3’ of the *nr2f1a* locus, with the conservation in 3’ regions of open chromatin extending to the syntenic gene *fam172a* [37] (**S2A Fig.**). We identified 7 conserved open regions flanking *nr2f1a* (**Fig. 1A-C**, **S2A,B,D,F,H Fig., and S3A-D Fig.**), which were subsequently analyzed via cloning into a GFP reporter vector [38]. Two of the putative enhancers did not drive significant GFP expression in transient transgenic embryos and were not studied further. The rest of the putative enhancers were named according to their 5’ and 3’ positions relative to *nr2f1a* (**Fig. 1A and S2A Fig.**). Those that did drive tissue-specific expression in transient transgenics were raised to confirm their expression in stable transgenic lines. Two of the transgenic enhancer reporters showed expression within subdomains that are reminiscent of the endogenous extra-cardiac *nr2f1a* expression [39]. *5’reg1-nr2f1a* was expressed in the anterior hindbrain and branchial arches (**S2C Fig.**). *3’reg4-nr2f1a* was expressed in the medial eye, nasal pits, and anterior branchial arch (**S2I Fig.**). Two putative enhancers, *3’reg2-nr2f1a* and *3’reg3-nr2f1a,* were expressed within the heart by 48 hpf, but did not show atrial-specific expression (**S2E,G Fig.**). However, the *3’reg1-nr2f1a* enhancer (henceforth referred to as *3’reg1*) showed expression specifically within ACs by 24 hpf, which was maintained through 72 hpf (**Fig. 1D-H**). Moreover, the strongest expression of the *3’reg1* reporter was at the venous pole of the atrium and it tapered toward the atrioventricular canal/outflow pole of the atrium (**Fig. 1E-H**). Although *nr2f1a* expression is first detectable by early somitogenesis stages in the ALPM [13], with the present analysis we did not identify CREs that drove expression during earlier stages in the ALPM. Thus, we identified a conserved putative *nr2f1a* enhancer that is able to promote expression within zebrafish ACs by the early heart tube stage.

**Figure 1.**
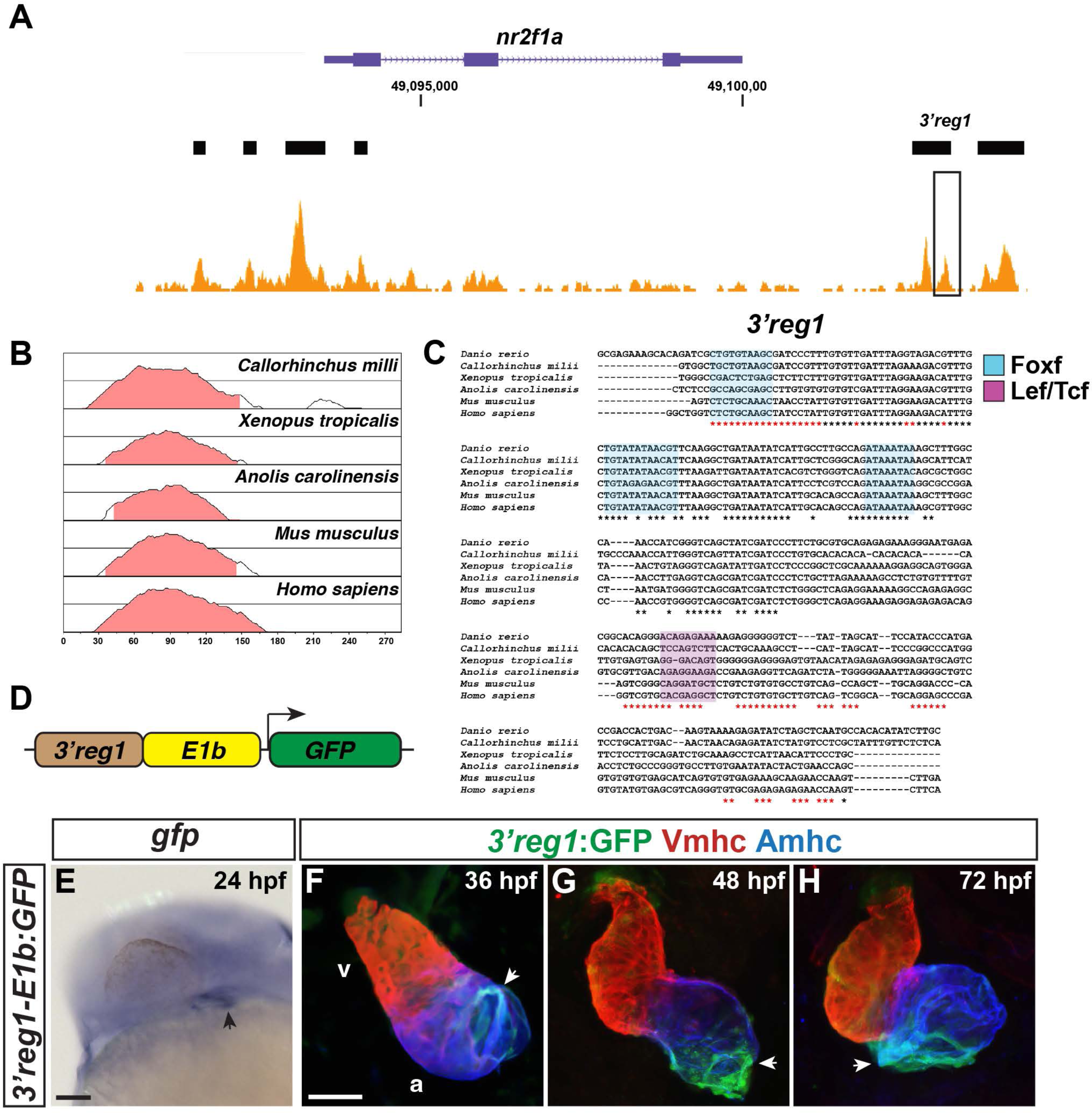
The *3’reg1-nr2f1a* enhancer is expressed in the atrium. **A)** ATAC-seq profiling from ACs showing regions of open chromatin near the *nr2f1a* locus (purple). Black bars indicate called regions of open chromatin. Black box indicates *3’reg1*. **B)** VISTA plot showing conservation of zebrafish *3’reg1* enhancer in *Callorhincus milli* (Australian ghostshark), *Xenopus tropicalis* (Tropical clawed frog), *Anolis carolinensis* (Green Anole), *Mus musculus* (House mouse), and *Homo sapiens* (human). Pink indicates >50% conservation of regulatory regions with zebrafish *3’reg1*. Median lines in individual VISTA plots indicate 75% conservation. **C)** Clustal alignment of *3’reg1* from the VISTA plot in indicated species. Black asterisks indicate conserved nucleotides. Red asterisks indicate partially conserved nucleotides. Conserved putative binding sites for Foxf (blue shade) and Lef/Tcf (purple shade) TFs. **D)** Schematic of the *3’reg1:GFP* reporter vector. **E)** *In situ* hybridization (ISH) for *gfp* in transgenic *3’reg1:GFP* embryo at 24 hpf (n=24). Venous pole of the heart tube (black arrowhead). View is lateral with anterior left and dorsal up. Scale bar: 200μm. **F-H)** Confocal images of hearts from transgenic *3’reg1:GFP* embryos stained for *3’reg1*:GFP (green), Vmhc (VCs – red), and Amhc (ACs – blue). 36 hpf (n=5), 48 hpf (n=7), 72 hpf (n=5). *3’reg1:GFP* in the atria of the hearts (white arrowhead). Images are frontal views with the arterial pole up. n indicates the number of embryos examined for a representative experiment. Scale bar: 50 μm. **** indicate P < 0.0001.

### Wnt signaling promotes Nr2f1a expression in ACs

To determine signals that could regulate *3’reg1*, we performed TF binding site (TFBS) analysis using the CIS-BP [40,41], TomTom [42,43], and JASPAR [44] databases. Among the different putative TFBSs identified within *3’reg1* were a nuclear hormone receptor (NHR) binding site, a Lef/Tcf binding site, and multiple Foxf binding sites (**Fig. 1C and S4A Fig.**), reflecting signals, including RA and Wnt signaling, that are involved in promoting the proper timing of AC differentiation from the posterior SHF progenitors in mice [15,16,32,45]. Furthermore, there was enriched conservation of nucleotides in the regions of the enhancers harboring the potential binding sites, although there was less conservation over the putative Lef/Tcf binding site (**Fig. 1C**). We next systematically interrogated the requirement of each of these sites in promoting *3’reg1:GFP* expression in ACs. Since RA signaling potentially regulates *Nr2f* expression during atrial differentiation [13,25,46], we initially examined the requirement of the NHR site (**S4A-C Fig.**). However, deletion of the NHR site did not alter expression of the reporter compared to the unaltered *3’reg1:GFP* reporter in transient transgenic embryos nor did modulation of RA signaling via treatment with RA and the RA signaling inhibitor DEAB affect expression in stable transgenic *3’reg1:GFP* lines (data not shown). Therefore, RA signaling does not appear to regulate *nr2f1a* expression through this enhancer.

Lef/Tcf TFs are mediators of canonical Wnt signaling [47,48], with numerous *in vivo* and *in vitro* studies having shown it controls differentiation of ACs [49,50]. Furthermore, we have shown that increased Wnt signaling during later somitogenesis immediately prior to the formation of the heart tube is necessary and sufficient to specifically produce an increase in ACs in zebrafish embryos [51]. Additionally, studies have demonstrated that Wnt signaling is expressed in a gradient extending from the venous pole of the zebrafish heart [51–53], which is reminiscent of the *3’reg1:GFP* reporter. We found that either deletion or targeted mutation of the Lef/Tcf site both resulted in an expansion of the reporter throughout the heart by 48 hpf relative to the *3’reg1:GFP* in transient transgenic embryos (**Fig. 2A-F**). As this result suggests the site may be used to restrict enhancer expression, we reasoned that Tcf7l1a (formerly called Tcf3a), a transcriptional repressor whose repression is relieved by Wnt signaling [54] that we have previously implicated in early cardiomyocyte development [53], may be limiting the expression of the reporter within the heart at these stages of cardiogenesis. To determine if Tcf71a limits expression of the reporter, *3’reg1:GFP* embryos were injected with an established morpholino (MO) that identically phenocopies the zebrafish *tcf7l1a* mutants [55]. We found that Tcf7l1a depletion produced pan-cardiac expression of the *3’reg1:GFP* reporter (**Fig. 2G-I**), supporting that Tcf7l1a is required to limit *3’reg1* reporter expression within the heart.

**Figure 2.**
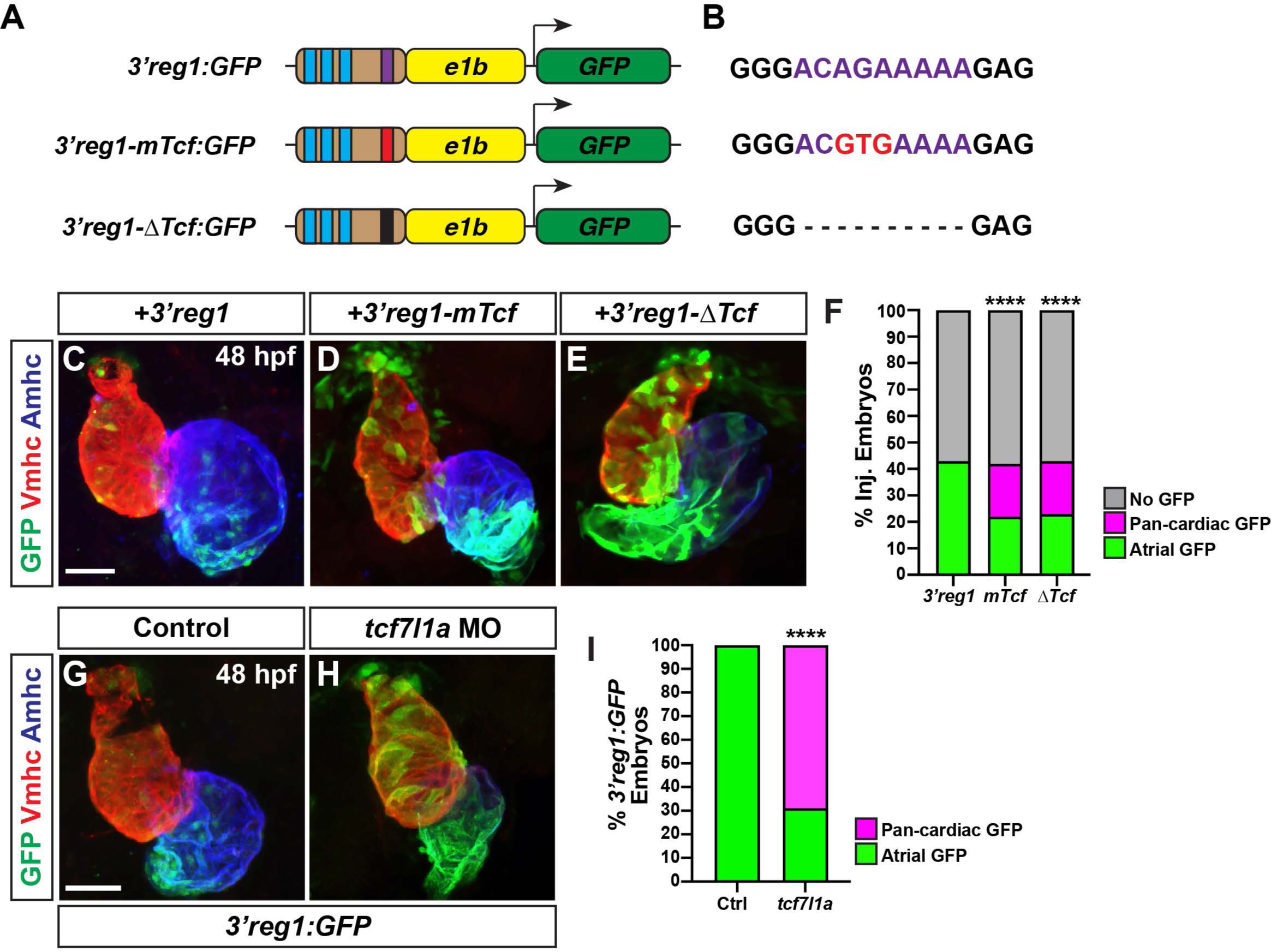
Tcf7l1a restricts *3’reg1* reporter expression within the heart. **A)** Schematics of *3’reg1:GFP* reporter constructs. Foxf sites (blue), Lef/Tcf site (purple), mutated Lef/Tcf site (red), and deleted Lef/Tcf site (black). **B)** Sequences of the WT (purple), mutated (red), and deleted Left/Tcf sites within the *3’reg1* enhancer. **C-E)** Confocal images of hearts from embryos injected with the *3’reg1:GFP*, *3’reg1-mTcf:GFP*, *3’reg1-ΔTcf:GFP* constructs. **F)** The percentage of transient transgenic embryos with reporter atrial, pan-cardiac, and lacking expression in their hearts. *3’reg1:GFP* (n=200); *3’reg1-mTcf:GFP* (n=101); *3’reg1-ΔTcf:GFP* (n=102). **G,H)** Confocal images of hearts from control and *tcf7l1a* MO-injected transgenic *3’reg1:GFP* embryos. **I)** The percentage of stable *3’reg1:GFP* embryos with reporter atrial and pan-cardiac expression in their hearts. Control (n=152); *tcf7l1a-MO* (n=92). Images are frontal views with the arterial pole up. Hearts are stained for *3’reg1*:GFP (green), Vmhc (red), Amhc (blue). Scale bars: 50 μm. **** indicate P < 0.0001.

As the activation of the transgenic *3’reg1:GFP* reporter via Tcf7l1a depletion would predict Wnt signaling should also regulate the *3’reg1:GFP* reporter expression within the heart, we manipulated Wnt signaling in *3’reg1:GFP* embryos at the 20 somite (s) stage with treatments of BIO and XAV939 (XAV), which respectively activate and inhibit Wnt signaling [56]. The 20s stage was chosen because we previously found that modulation of Wnt signaling could specifically affect AC number slightly prior to this stage at ∼16s [51,53]. Although the effect of Wnt signaling on AC differentiation was not examined previously at the 20s stage, this stage was also just prior to when we can detect transgenic *3’reg1:GFP* reporter expression. We found that by 48 hpf the hearts of the embryos treated with BIO showed pan-cardiac GFP expression (**Fig. 3A,B,D**), similar to perturbation of the Lef/Tcf sites in the *3’reg1:GFP* reporter (**Fig. 2C-E**) and Tcf7l1a depletion (**Fig. 2G,H**), while XAV abrogated GFP expression within the atria (**Fig. 3A,C,D**). Since our data supported that Wnt signaling activates the *3’reg1:GFP* reporter, likely via derepression of Tcf7l1a [53], we tested if Wnt signaling at these stages impacts Nr2f1a expression within the heart. As a proxy for affecting Nr2f1a expression, we quantified the number of Nr2f1a^+^ cardiomyocyte nuclei within the hearts of 48 hpf embryos treated with BIO and XAV beginning at the 20s stage (**Fig. 3E-H**). Although we did not observe obvious ectopic Nr2f1a expression within ventricular cardiomyocytes (VCs) as with the *3’reg1:GFP* reporter, BIO treatment at the 20s stage produced a surplus of Nr2f1a^+^/Ahmc^+^ cardiomyocytes (ACs) by 48 hpf. Conversely, XAV treatment caused a decrease in Nr2f1a^+^/Amhc^+^ cardiomyocytes (ACs) within the atrium, with the Amhc^+^ ACs qualitatively having a reduction in Nr2f1a expression within their nuclei. Based on the effects of Wnt signaling on Nr2f1a in ACs and our previous observations that manipulation of Wnt signaling at similar somitogenesis stages specifically affected AC production [51], we tested the impact of Wnt manipulation on ACs using the *myl7:DsRed2-NLS* transgene [57]. In agreement with the assessment of Nr2f1a^+^ cardiomyocytes, BIO and XAV treatment at the 20s stage respectively produced a surplus and decrease in the number of ACs (*myl7:*DsRed2-NLS^+^/Amhc^+^ cardiomyocytes) by 48 hpf (**Fig. 3I-L**). However, we did not detect changes in the number of VCs (**S5A Fig.**), similar to what we have reported previously [51,53]. Thus, Wnt signaling is necessary and sufficient at the 20s stage to activate the transgenic *3’reg1:GFP* reporter and promote a surplus of differentiated ACs, likely through derepression of Tcf7l1a.

**Figure 3.**
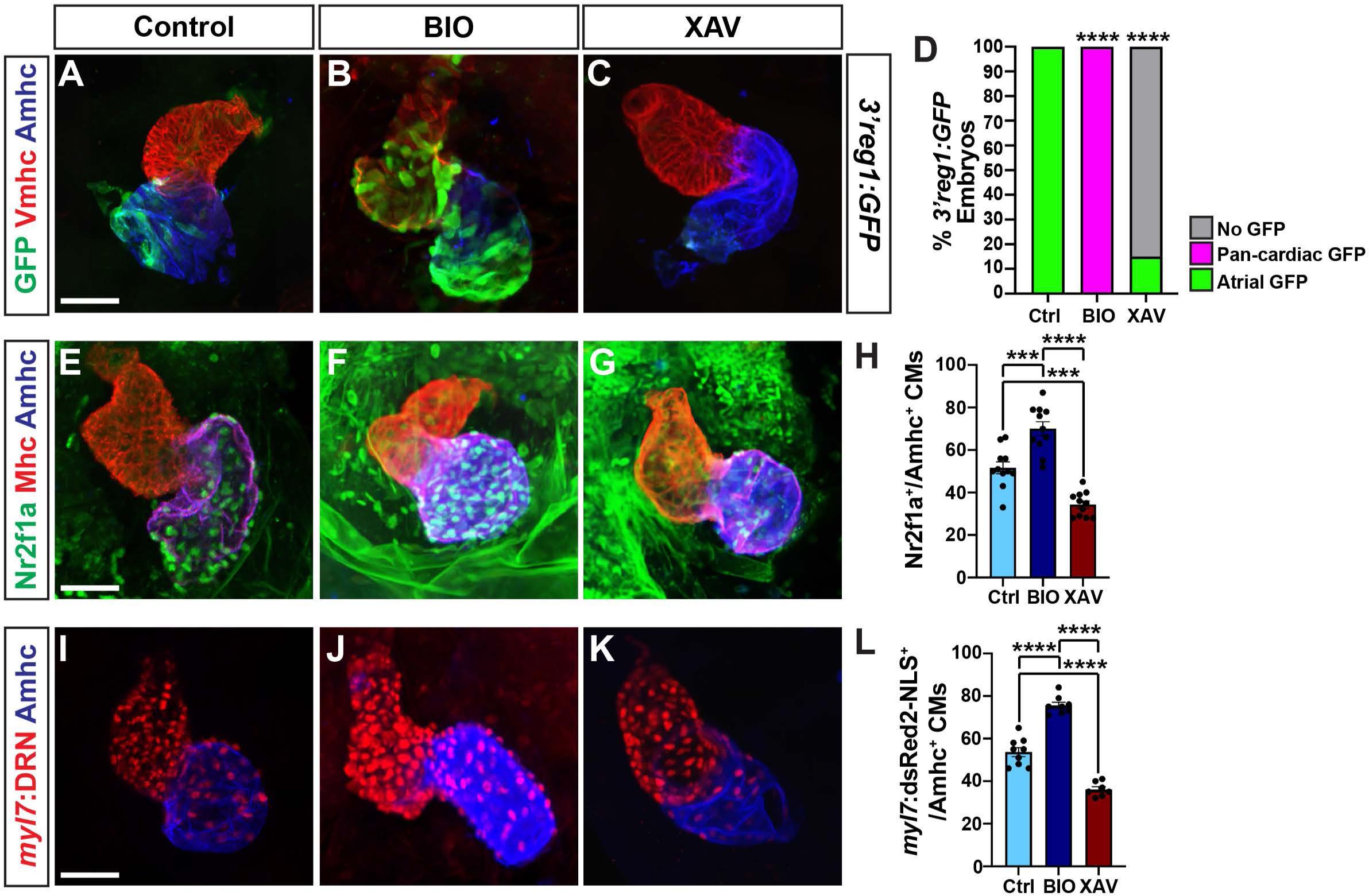
Wnt signaling promotes *3’reg1* reporter expression and AC differentiation. **A-C)** Confocal images of hearts from control, BIO-, and XAV-treated *3’reg1:GFP* embryos stained for *3’reg1*:GFP (green), Vmhc (red), Amhc (blue). **D)** The percentage of control and treated *3’reg1:*GFP embryos with atrial, pan-cardiac, and lacking expression in their hearts. Control (n=35); BIO (n=32); XAV (n=49). **E-G)** Confocal images of hearts from control, BIO-, and XAV-treated embryos stained for Nr2f1a (green), Mhc (pan-cardiac, red), Amhc (blue). **H)** The number of Nr2f1a^+^/Amhc^+^ cardiomyocytes with the hearts of control, BIO-, and XAV-treated embryos. Control (n=12); BIO (n=11); XAV (n=11). **I-K)** Confocal images of hearts from control, BIO-, and XAV-treated *myl7:DsRed2-NLS* (*myl7:DRN*) embryos stained for DsRed-NLS (pan-cardiac nuclei - red) and Amhc (blue). **L)** The number of *myl7:*DsRed2-NLS^+^/Amhc^+^ cardiomyocytes (ACs) within the hearts of control, BIO-, and XAV-treated *myl7:DsRed2-NLS* embryos. Control (n=9); BIO (n=10); XAV (n=8). Scale bars: 50 μm. Error bars in graphs indicate s.e.m.. *** indicate P < 0.001. **** indicate P < 0.0001.

### Foxf1 promotes Nr2f1a expression in ACs

In mice, Foxf1 and Foxf2 TFs control the proper differentiation of posterior SHF progenitors downstream of Hh signaling [30,58]. Thus, we were intrigued by the conserved Foxf sites within the *3’reg1* enhancer (**Fig. 1C**). We found that in transient transgenic embryos deletion or mutation of each of the 3 Foxf binding sites diminished the percentage of embryos with GFP in their atria relative to injection of the wild-type (WT) *3’reg1* reporter (**Fig. 4A-C**), suggesting the putative Foxf sites may be required to promote *3’reg1:GFP* expression. To further test this hypothesis, murine *Foxf1* mRNA was injected into *3’reg1:GFP* transgenic embryos. We found Foxf1 was sufficient to expand and promote *3’reg1:GFP* expression into VCs by 48 hpf (**Fig. 4D,E,G**). Conversely, injecting a dominant-negative murine *Foxf1* (*dnFoxf1*) mRNA into the *3’reg1:GFP* transgenic embryos was able to inhibit reporter expression within the heart (**Fig. 4D,F,G**). Although zebrafish mutants for *foxf1* have been described [59], if they have cardiac defects was not reported. Nevertheless, we sought to determine if *foxf1* is required for *3’reg1:GFP* reporter expression and we targeted zebrafish *foxf1* using the CRISPR-Cas12a system [60–62] with guides we found were highly efficient in creating deletions (**S6A,B Fig.**). Although we did not observe overt cardiac defects, injection of the *foxf1* CRISPRs was sufficient to reduce expression of the *3’reg1:GFP* reporter in transgenic embryos (**S6C-E Fig.**). Given the effects on the *3’reg1:GFP* reporter, we asked if Foxf1 was sufficient to promote Nr2f1a^+^ ACs and increase the number of differentiated ACs. We found that embryos injected with *Foxf1* or *dnFoxf1* mRNA respectively showed increases and decreases in both Nr2f1a^+^/Amhc^+^ and *myl7:DsRed2-NLS^+^/*Amhc*^+^*cardiomyocytes within their hearts at 48 hpf (**Fig. 4H-O**), without affecting the number of VCs (**S5B Fig.**). As with Wnt inhibition, embryos injected with the *dnFoxf1* mRNA qualitatively had reduced expression of Nr2f1a within the atrium at 48 hpf (**Fig. 4J**). Thus, our data suggest that Foxf1 can activate the transgenic *3’reg1:GFP* reporter and promote an increase in differentiated Nr2f1a-expressing ACs.

**Figure 4.**
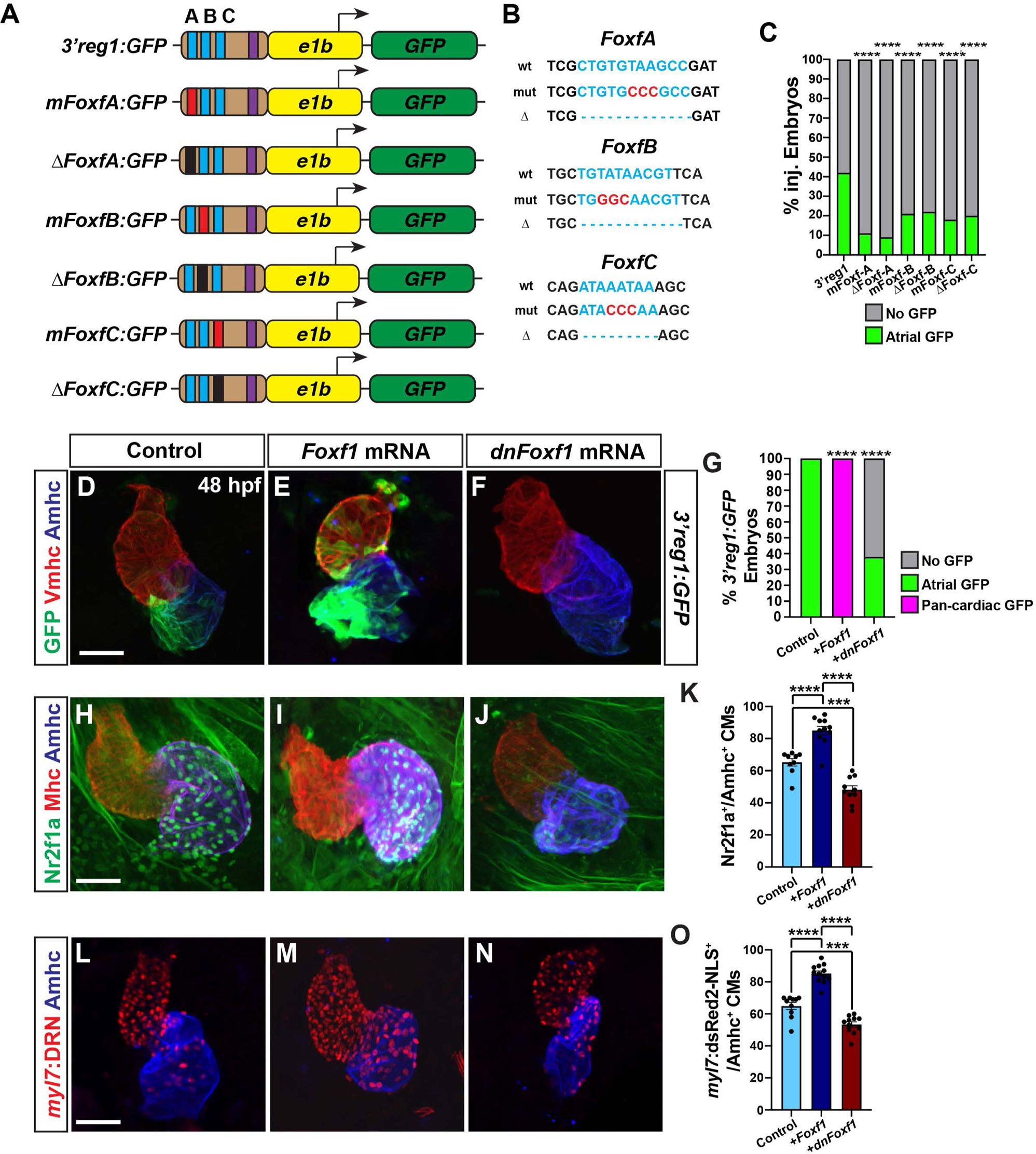
Foxf1 promotes *3’reg1* reporter expression within the heart. **A)** Schematics of *3’reg1:GFP* reporter constructs with WT, deleted, and mutated Foxf sites. Foxf sites (blue), mutated Foxf sites (red), deleted Foxf sites (black), and Lef/Tcf site (purple). **B)** Sequences of the WT (blue), mutated (red), and deleted (dashes) Foxf sites within the *3’reg1* enhancer. **C)** The percentage of transient transgenic embryos with reporter expression in the atria. Control (*3’reg1*) (n=150); *mFoxf-A* (n=60); *ΔFoxf-A* (n=101); *mFoxf-B* (n=49); *ΔFoxf-B* (n=200); *mFoxf-C* (n=98); *ΔFoxf-C* (n=195). **D-F)** Confocal images of hearts from control uninjected, *Foxf1* mRNA and *dnFoxf1* mRNA injected *3’reg1:GFP* embryos stained for *3’reg1*:GFP (green), Vmhc (red), Amhc (blue). **G)** The percentage of injected *3’reg1:GFP* embryos with atrial expression, pan-cardiac expression, and lacking expression. Control (n=35); *Foxf1* (n=29); *dnFoxf1* (n=59). **H-J)** Confocal images of hearts from control, *Foxf1* mRNA, and *dnFoxf1* mRNA injected embryos stained for Nr2f1a (green), Mhc (red), Amhc (blue). **K)** The number of Nr2f1a^+^/Amhc^+^ cardiomyocytes (ACs) within the hearts of control, *Foxf1* mRNA, and *dnFoxf1* injected embryos. Control (n=8); *Foxf1* (n=11); *dnFoxf1* (n=8). **L-N)** Confocal images of hearts from control, *Foxf1* mRNA, and *dnFoxf1* injected *myl7:DsRed2-NLS* (*myl7:DRN*) embryos stained for DsRed-NLS (red) and Amhc (blue). **O)** Quantification of the number of *myl7:*DsRed2-NLS^+^/Amhc^+^ cardiomyocytes (ACs) within the hearts of control, *Foxf1* mRNA, and *dnFoxf1* injected *myl7:DsRed2-NLS* embryos. Control (n=10); *Foxf1* (n=13); *dnFoxf1* (n=11). Scale bars: 50 μm. Error bars in graphs indicate s.e.m.. *** indicate P < 0.001. **** indicate P < 0.0001.

### Hh signaling represses *3’reg1* reporter expression

In mice, Hh signaling has been established as a key regulator of posterior SHF differentiation upstream of factors, including Foxf1 and Foxf2, and Wnt signaling [29–31], and has previously been implicated in regulating Nr2f2 expression in P19 cells [63]. Thus, we asked if Hh signaling could affect the *3’reg1:GFP* reporter within the atria. Transgenic *3’reg1:GFP* embryos were treated with the Hh signaling inhibitor Cyclopamine (CYA) [64] and the Hh signaling activator SAG [65] beginning at the 20s stage, which is after previous reports demonstrated that Hh signaling affects the production of cardiac progenitors [55]. However, in contrast to the effects of Wnt and Foxf1 manipulation, embryos treated with CYA at the 20s stage unexpectedly showed a modest expansion of the *3’reg1:GFP* reporter within the atrium, while SAG-treated *3’reg1:GFP* embryos showed a reduction of *3’reg1*:GFP-expression (**S7A-D Fig.**). To confirm that Hh signaling represses *3’reg1:GFP* reporter expression, we examined the *3’reg1:GFP* reporter in *smoothened* (*smo)* mutants. Consistent with CYA treatments, we found a modest expansion of the *3’reg1:GFP* reporter within the atria of *smo* mutants (**S7E-G Fig.**). However, despite the effects on the *3’reg1:GFP* reporter within the atrium, modulation of Hh signaling did not affect the number of Nr2f1a^+^ ACs (**S7H-K Fig.**). As Hh signaling may be upstream of Wnt signaling in posterior SHF differentiation in mammals [31,66], we also concurrently manipulated Wnt and Hh signaling at the 20s stage with the aforementioned drugs. We found that simultaneously inhibiting or activating Wnt and Hh signaling on decreased and increased expression of the *3’reg1:GFP* reporter within the heart, similar to inhibiting or activation of Wnt signaling alone (**S7L-O Fig.**). Therefore, our data suggest that in contrast to mammalian venous pole differentiation Hh signaling is likely not affecting AC differentiation at these stages of cardiogenesis nor is it promoting Nr2f expression in ACs upstream of Wnt or Foxf1 in zebrafish embryos.

### Wnt signaling functions downstream of Foxf1

To decipher the functional relationship between Wnt signaling/Tcf7l1a and Foxf1 on activation of the *3’reg1:GFP* reporter within the atrium, we simultaneously manipulated these signals within the transgenic embryos. While *3’reg1:GFP* embryos injected with *Foxf1* mRNA had pan-cardiac expression, the majority of *Foxf1* mRNA-injected *3’reg1:GFP* embryos treated with XAV at the 20s stage lose *3’reg1:GFP* reporter expression within the atrium by 48 hpf, similar to XAV treatments alone (**Fig. 5A-E**). We next simultaneously injected *Foxf1* and zebrafish *tcf7l1a* mRNAs into the *3’reg1:GFP* embryos. Consistent with the Tcf7l1a functioning as a repressor, *tcf7l1a* overexpression abrogated *3’reg1:GFP* reporter expression within the atrium (**Fig. 5F,H,J**). When both TFs were co-overexpressed in *3’reg1:GFP* embryos, we did not observe pan-cardiac expression of the *3’reg1:GFP* reporter within the atrium at 48 hpf (**Fig. 5F-J**). Next, we co-injected the *tcf7l1a* MO and *dnFoxf1* mRNA into the *3’reg1:GFP* embryos. In this scenario, we found that the *dnFoxf1* was not sufficient to block the pan-cardiac expression induced by Tcf7l1a depletion (**Fig. 5K-O**). Since these data suggested that Wnt signaling may function downstream of Foxf1, we asked if Foxf1 was sufficient to promote an increase in Wnt signaling within embryos. RT-qPCR on 24 hpf embryos injected with *Foxf1* mRNA promoted a modest but significant increase in the expression of *axin1*, a Wnt responsive gene that negatively regulates canonical Wnt signaling [67], and a decrease in the expression of the Wnt signaling inhibitor *dkk1a* [67] (**S8A,B Fig.**). Cumulatively, these data suggest that Wnt signaling functions downstream, or parallel, to Foxf1 in the activation of the *3’reg1:GFP* reporter.

**Figure 5.**
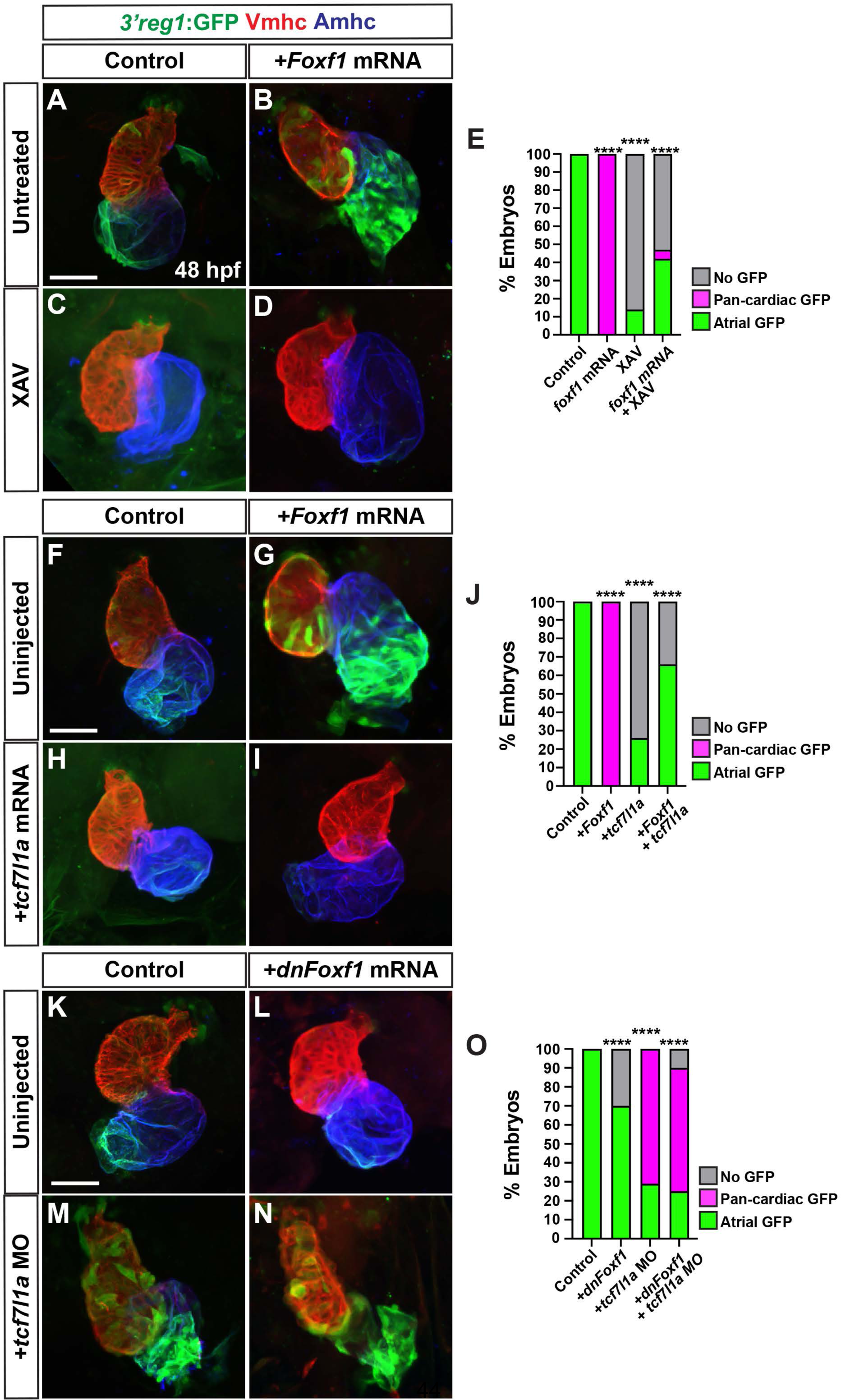
Tcf7l1a limits the ability of Foxf1 to promote *3’reg1* reporter expression. **A-D)** Confocal images of hearts from control, *Foxf1* mRNA injected, XAV-treated, and *Foxf1* mRNA injected plus XAV-treated *3’reg1:GFP* embryos. **E)** The percentage of control, *Foxf1* mRNA injected, XAV-treated, and *Foxf1* mRNA injected plus XAV-treated *3’reg1:GFP* embryos with atrial expression, pan-cardiac expression, and lacking expression. Control (n=26); *Foxf1* (n=24); XAV n=98; *Foxf1*-XAV (n=103). **F-I)** Confocal images of hearts from control, *Foxf1* mRNA injected, *tcf7l1a* mRNA injected, and *Foxf1* mRNA and t*cf7l1a* mRNA co-injected *3’reg1:GFP* embryos. **E)** The percentage of control, *Foxf1* mRNA injected, *tcf7l1a* mRNA injected, and *Foxf1* mRNA and t*cf7l1a* mRNA co-injected *3’reg1:GFP* embryos with atrial expression, pan-cardiac expression, and lacking expression. Control (n=39), *Foxf1* (n=34); *tcf7l1a* (n=95); *Foxf1*-*tcf7l1a* (n=51). **K-N)** Confocal images of hearts from control, *dnFoxf1* mRNA injected, *tcf7l1a* MO injected, and *Foxf1* mRNA and t*cf7l1a* MO co-injected *3’reg1:GFP* embryos. **O)** The percentage of control, *dnFoxf1* mRNA injected, *tcf7l1a* MO injected, and *Foxf1* mRNA and t*cf7l1a* MO *3’reg1:*GFP embryos with atrial expression, pan-cardiac expression, and lacking expression. Control (n=27), *dnFoxf1* (n=102); *Tcf7l1a-MO* (n=96); *dnFoxf1-tcf7l1aMO* (n=100). Hearts are stained for *3’reg1*:GFP (green), Vmhc (red), Amhc (blue)Scale bars: 50 μm. **** indicate P < 0.0001.

### Wnt signaling and Foxf1 require Nr2f1a to promote surplus ACs

Nr2f1a is required for AC differentiation within the early zebrafish heart [12]. As our results support that enhanced Wnt signaling and Foxf1 are sufficient to promote more differentiated ACs, we asked if this ability was dependent on Nr2f1a. Embryos resulting from crosses of *nr2f1a*^+/−^ fish that carry the *myl7:DsRed2-NLS* transgene were treated with BIO at the 20s stage. We found that BIO treatment was not sufficient to produce an increase of ACs in 48 hpf *nr2f1a* mutants, while it was sufficient to produce an increase in the number of ACs in their WT siblings (**Fig. 6A-E**). Similarly, injection of *Foxf1* mRNA into embryos resulting from crosses of *nr2f1a*^+/−^ fish that carry the *myl7:DsRed2-NLS* transgene showed that *Foxf1* mRNA was no longer sufficient to produce a surplus of ACs in *nr2f1a* mutants, unlike their control sibling embryos (**Fig. 6F-J**). Thus, our data support that Nr2f1a is required for Wnt signaling and Foxf1 to be sufficient to promote a surplus of ACs.

**Figure 6.**
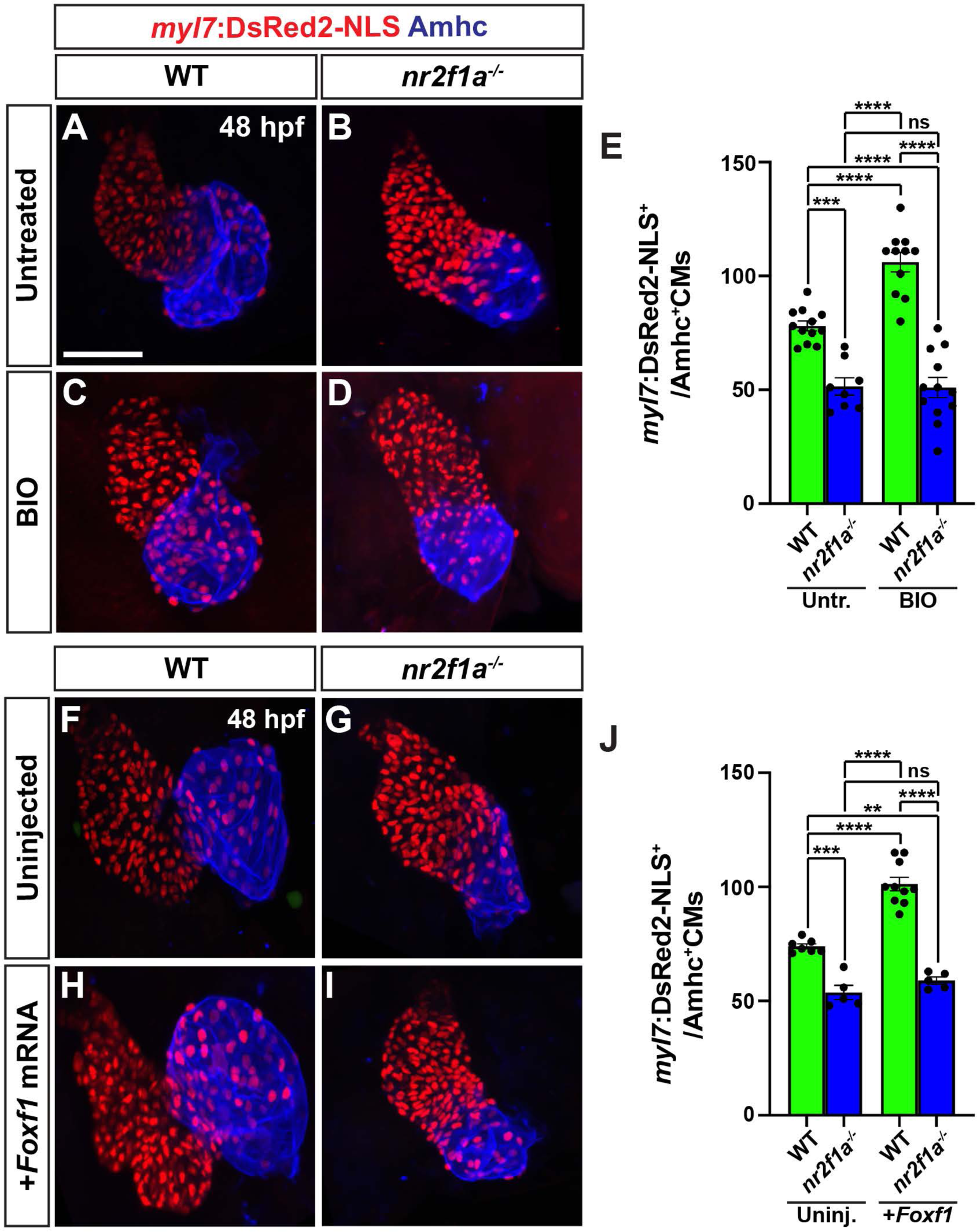
Wnt signaling and Foxf1 require Nr2f1a to promote a surplus of cardiomyocytes. **A-D)** Confocal images of hearts from untreated and BIO-treated WT and *nr2f1a*^−/−^ *myl7:DsRed2-NLS* embryos. **E)** The number of *myl7:*DsRed2-NLS^+^/Amhc^+^ cardiomyocytes in hearts of untreated and BIO-treated WT and *nr2f1a*^−/−^ *myl7:DsRed2-NLS* embryos. Untreated WT (n=12); untreated *nr2f1a*^−/−^ (n=7); treated WT (n=11); treated *nr2f1a*^−/−^ (n= 10). **F-I)** Confocal images of hearts from uninjected and *Foxf1* mRNA-injected WT and *nr2f1a*^−/−^ *myl7:DsRed2-NLS* embryos. **J)** The number of *myl7:*DsRed2-NLS^+^/Amhc^+^ cardiomyocytes in uninjected and *Foxf1* mRNA-injected WT and *nr2f1a*^−/−^ *myl7:DsRed2-NLS* embryos. Uninjected WT (n=7); Uninjected *nr2f1a*^−/−^ (n=5); Inj. WT (n=10); Injected *nr2f1a*^−/−^ (n=5). Hearts are stained for DsRed-NLS (red) and Amhc (blue). Scale bars: 50 μm. Error bars in graphs indicate s.e.m.. ** indicate P < 0.01. **** indicate P < 0.001. **** indicate P < 0.0001. ns indicates not a statistically significant difference.

### Wnt signaling and Foxf1 require *3’reg1* to promote ACs

Since the *3’reg1:GFP* reporter is expressed in ACs, we wanted to determine if the endogenous *3’reg1* enhancer is required *in vivo* to promote or maintain Nr2f1a expression in ACs. Therefore, we deleted the endogenous *3’reg1* enhancer in zebrafish using a CRISPR-Cas12a system [68] (**Fig. 7A and S9A,B Fig.**). We found that embryos with homozygous deletion of the endogenous *3’reg1* enhancer overtly did not show defects (**S9C,D Fig.**). Furthermore, there was no difference in the number of ACs (Nr2f1a^+^/Amhc^+^ cardiomyocytes) in the hearts of *3’reg1* deletion mutants compared to their WT sibling embryos at 48 hpf (**Fig. 7B,C,F,G,H,K**), suggesting that *in vivo* deletion of the *3’reg1* enhancer alone is not sufficient to abrogate proper N2f1a expression within the atrium. Consequently, we then asked if Wnt signaling and Foxf1 require the endogenous *3’reg1* enhancer to be able to produce a surplus of Nr2f1a^+^ ACs. We found that both BIO treatment and Foxf1 mRNA injection were unable to produce a surplus of Nr2f1a^+^/Amhc^+^ cardiomyocytes within the hearts of homozygous *3’reg1* deletion mutants (**Fig. 7B-K**). Overall, these data suggest that the endogenous *3’reg1* is required to mediate the sensitivity of Nr2f1a and AC differentiation to Wnt signaling and Foxf1 in the zebrafish atrium.

**Figure 7.**
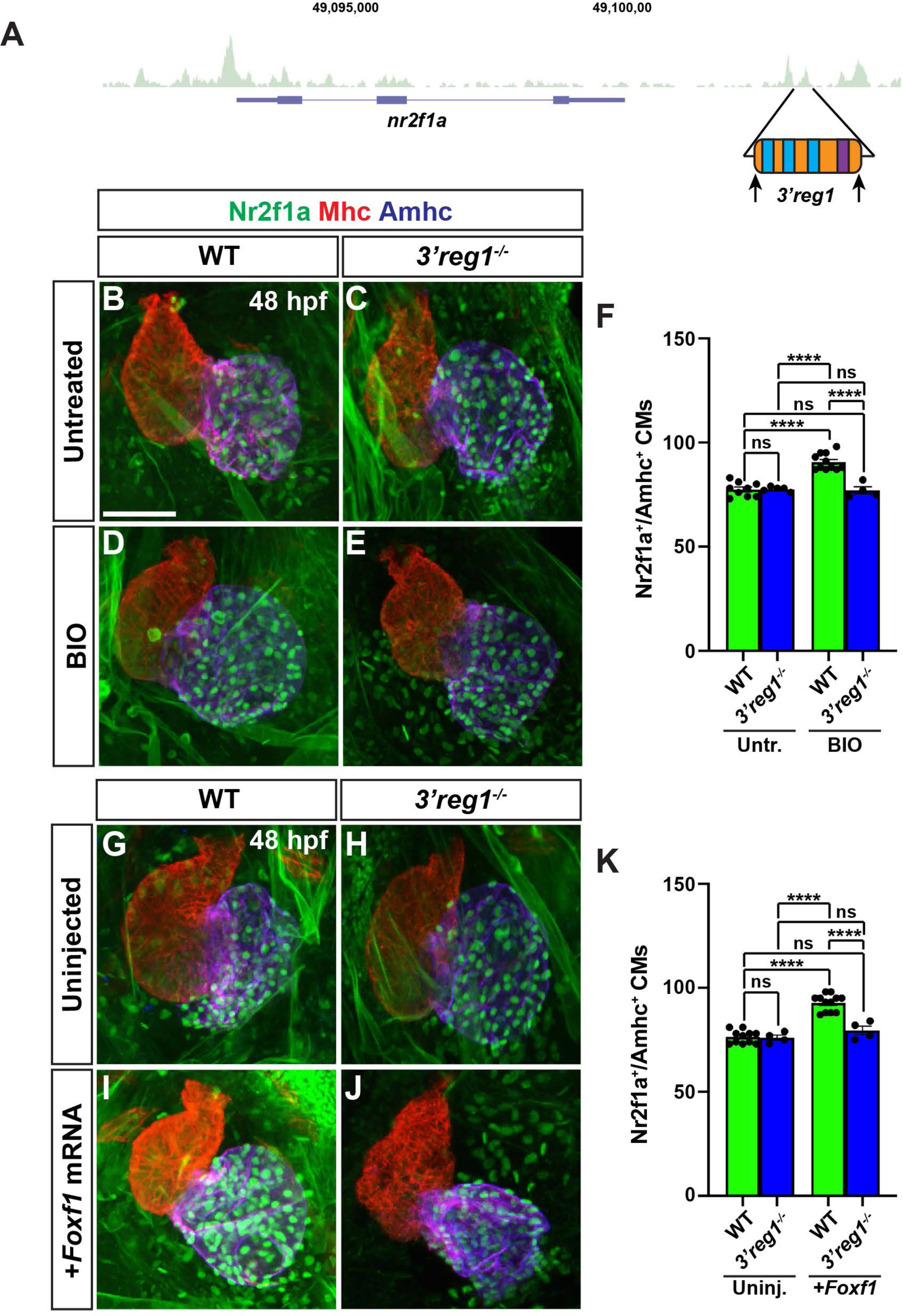
Wnt signaling and Foxf1 require 3*’reg1* to promote a surplus ACs. **A)** Schematic of *3’reg1* enhancer deletion generated using CRISPR/Cas12a. Arrows indicate location of the guides. **B-E)** Confocal images of hearts from untreated and BIO-treated WT and *3’reg1^Δ/Δ^* embryos stained for Nr2f1a (green), Mhc (red), and Amhc (blue). BIO was unable to promote an increase in ACs in *3’reg1^−/−^* embryos. **E)** The number of Nr2f1a^+^/Amhc^+^ cardiomyocytes in untreated and BIO-treated WT and *3’reg1^−/−^* embryos. Untreated WT (n=9); untreated *3’reg1^−/−^* (n=5); treated WT (n=11); treated *3’reg1^−/−^* (n=4). **F-I)** Confocal images of hearts from uninjected and *Foxf1* mRNA-injected WT and *3’reg1^−/−^*embryos stained for Nr2f1a (green), Mhc (red), and Amhc (blue). **J)** The number of Nr2f1a^+^/Amhc^+^ cardiomyocytes in uninjected and *Foxf1* mRNA-injected WT and Nr2f1a^+^ *3’reg1^−/−^* embryos. Uninjected WT (n=11); Uninjected *3’reg1^−/−^* (n=4); Injected WT (n=12); Injected *3’reg1^−/−^* (n= 4). Scale bar: 50 μm. Error bars in graphs indicate s.e.m.. **** indicate P < 0.0001. ns indicates not a statistically significant difference.

## Discussion

In the present study, our data show that in zebrafish embryos Wnt signaling and Foxf1 function upstream of Nr2f1a in promoting AC differentiation via regulating a conserved *Nr2f1* enhancer, which is located ∼2.5kb 3’ to zebrafish *nr2f1a* (**Fig. 8**). Previous work has placed both these signals downstream of Hh signaling in a regulatory network that controls the differentiation of ACs from the posterior SHF in mice [31,32,66]. Thus, our data support that there are conserved roles for these downstream factors in promoting the proper number of differentiated ACs in the hearts of vertebrates. Numerous studies have investigated the different roles of canonical Wnt signaling in cardiac development in mice, zebrafish, and stem cells [32,33,47,53,69]. Furthermore, our previous study examining the temporal requirements of Wnt signaling in zebrafish showed that it is necessary and sufficient to specifically promote surplus ACs during later somitogenesis immediately prior to the formation of the heart tube [51,53], a time point when posterior SHF progenitors are adding to the zebrafish heart [70]. Additional work in zebrafish has shown that Wnt signaling at the lateral borders of the heart field and at the venous pole is required for the differentiation of pacemaker cardiomyocytes [71]. Our present study corroborates the role for Wnt signaling in promoting the proper number of ACs as the heart tube is forming during late somitogenesis, which we propose is through affecting the differentiation of ACs from the zebrafish posterior SHF. Furthermore, our work supports that the ability of Wnt signaling to promote ACs necessitates Nr2f1a, whose expression is regulated via the derepression of Tcf7l1a.

**Figure 8.**
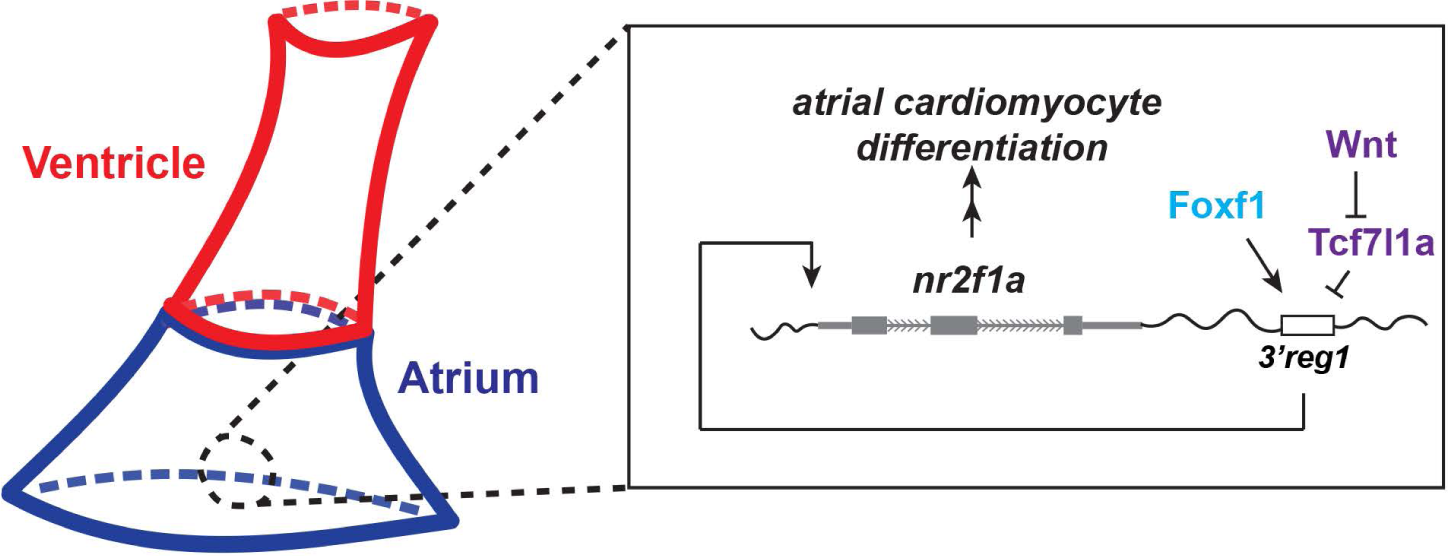
A Wnt-Foxf1-Nr2f1a cascade directs atrial cardiomyocyte differentiation. Model depicting the Wnt signaling and Foxf1-dependent regulatory network that controls Nr2f1a expression and the role of the *3’reg1* enhancer in the atrial cardiomyocyte differentiation during early heart tube development.

Despite the established roles of Foxf1/2 TFs in regulating differentiation of the posterior SHF in mice [58,72,73], if Foxf TFs play similar roles in zebrafish heart development has not been investigated. Previous work examining Foxf factors in zebrafish demonstrated redundant requirements for Foxf1/2 TFs in craniofacial development [59,74]. Similar to the roles of Foxf1/2 in mice [30,58,72], our data suggest that the Foxf TFs are necessary and sufficient to promote the formation of ACs in zebrafish embryos. Although our data did not yet specifically address the temporal or spatial requirements of Foxf factors, the necessity of Nr2f1a in promoting the proper number of differentiated ACs would also support a model that affects the differentiation of zebrafish posterior SHF progenitors. Thus, as with Wnt signaling, our data support a conserved requirement for Foxf TFs upstream of Nr2f1a in controlling the differentiation of ACs at the venous pole in vertebrates. However, although we found that depletion of *foxf1* affected the expression of the *3’reg1:GFP* reporter, zebrafish *foxf1* crispants did not overtly have cardiac defects. Similarly, Wnt signaling and Foxf1 could induce ectopic *3’reg1:GFP* reporter in VCs, but neither were able to induce ectopic Nr2f1a in VCs. Hence, the reporter may be more sensitive to manipulation of Wnt signaling and Foxf factors than endogenous Nr2f1a due to a lack of input from additional CREs. Nevertheless, we postulate that multiple *foxf* factors in zebrafish likely function redundantly to promote AC differentiation of zebrafish, as with Foxf1/2 in mice [58,72], which will be investigated more in the future.

Integrating how Wnt signaling and Foxf TFs function upstream of Nr2f1a, our analysis supports that Wnt signaling, via derepression of Tcf7l1a, functions downstream, or at the level, of Foxf1. This functional relationship is reminiscent of other developmental contexts, such as osteogenesis [75,76], where Foxf factors have been found to regulate the canonical Wnt signaling pathway [77]. Despite their functional relationship in promoting the differentiation of zebrafish ACs, we have not determined the specific mechanism by which Foxf factors regulate Wnt signaling in this context. Given the adjacent Foxf sites and Lef/Tcf site within the *3’reg1* CRE, one might predict that Foxf factors, which can act as pioneer factors [78,79], facilitate the derepression of Tcf7l1a via making the CRE accessible to Wnt signaling. However, our data would argue against this model, as we find Tcf7l1a does not appear to require Foxf for the derepression. Another possibility is that Foxf reinforces the activation of the CRE via simultaneously acting on the *3’reg1* enhancer and independently promoting Wnt signaling via derepression of Tcf7l1a. Additionally, while Foxf1 can function as a transcriptional activator [80], it is predominantly known as a transcriptional suppressor [81,82]. The dependence of the *3’reg1:GFP* reporter on the putative Foxf sites would minimally suggest that in this context these sites are being utilized by a transcriptional activator, which correlates with the evidence that Foxf1 is able to promote *3’reg1:GFP* reporter expression and atrial differentiation. Nevertheless, we acknowledge that in the absence of evidence for direct binding to the endogenous *3’reg1* enhancer, other scenarios are plausible, including that Foxf1 represses a Wnt inhibitor. However, in mice, in addition to repression of factors that limit differentiation of the posterior SHF, Foxf1/2 are also required for the activation of chamber-specific differentiation programs [30] which could also suggest that it has dual functions as a repressor and activator in this context.

In mammals, Hh signaling sits atop a regulatory network that controls the proper timing of differentiation from posterior SHF progenitors through Tbx5 and Foxf TFs, and Wnt and RA signaling [30,32,69]. Although our data support that downstream factors in this regulatory network, Wnt signaling and Foxf TFs, are conserved in regulating posterior SHF addition in zebrafish, our data and previous studies do not support the conservation of the upstream factors [30,31]. Previous work established that in zebrafish Hh signaling is required during early patterning stages to promote the proper number of differentiated ACs and VCs [55], while a requirement at later stages of somitogenesis and posterior SHF addition equivalent to those examined in this study was not found. Consistent with the previous observations of the temporal requirements of Hh signaling [55,83], we did not find it was required upstream of Wnt signaling or Foxf. Instead, we found that there was a minimal effect of Hh signaling repressing the *3’reg1:GFP* reporter, which was the opposite of what would be expected from the mammalian data. Furthermore, we did not find evidence that Hh regulates Nr2f1a or affects the number of differentiated ACs in zebrafish during late somitogenesis and early heart tube formation. Previous work has also established that Tbx5, which functions downstream of Hh and RA signaling in mice in posterior SHF differentiation [30–32], is not required for zebrafish atrial differentiation [84–86]. It is interesting to consider the lack of conservation of these upstream factors in the regulation of the vertebrate posterior SHF. The posterior SHF-derived cells that contribute to the single atrium and venous pole in zebrafish are significantly less than that found in mice [70,83]. By virtue of its lack of requirement, our work would support the proposed hypothesis whereby heterochrony of a Hh-dependent regulatory network may have contributed to the enlargement of the posterior SHF and the concurrent the evolution of the pulmonary system in air-breathing vertebrates [69,83].

Specifically evaluating the *3’reg1* enhancer, our data support that it is highly conserved among vertebrates, but is absent in agnathans (hagfish and lamprey) and birds. A lack of this CRE in agnathans could suggest that it evolved beginning in cartilaginous fishes. Although not explored here, *Nr2f2* genes, which diverged from *Nr2f1* early in the vertebrate lineage [37], also have a conserved putative enhancer located in a similar region (**S10 Fig.**), suggesting these enhancers may have had a similar origin at least in agnathans prior to the divergence of these paralogs. Although the *Nr2f1 3’reg1* is lost in birds, the conservation of the putative *Nr2f2-3’reg1* within birds could reflect the greater reliance of *Nr2f2* within their hearts similar to its role in mammals. The general conservation of the *3’reg1* in vertebrates is interesting in light that it has been suggested that CREs regulating cardiac expression are actually poorly conserved in vertebrates [87]. Additionally, deletion of the *3’reg1* did not cause overt loss of or failure to maintain Nr2f1a and ACs deficits. Thus, other CREs are likely involved in or compensate for the loss of the *3’reg1*, which is not uncommon [88], and may also be a reason its loss could be tolerated in agnathans and birds. We focused on the *3’reg1* because of the functional importance of *nr2f1a* in zebrafish atrial differentiation. Our data support that Wnt signaling and Foxf factors require the endogenous *3’reg1* enhancer to be able to augment the number of ACs, even if the element itself is not required to promote or maintain normal Nr2f1a expression within the heart. Furthermore, although this was the only conserved *Nr2f1* CRE thus far that we identified promotes AC-specific expression, we did find nearby CREs that promoted expression more broadly in cardiomyocytes. We have not yet examined the coordination of these CREs and if expression driven with the region 3’ to *nr2f1a* encompassing all the CREs better reflects endogenous *nr2f1a* within ACs. Additionally, as the transgenic *3’reg1* reporter expression initiates at early heart tube stages, which is later than endogenous *nr2f1a* expression occurs in the anterior lateral plate mesoderm during early somitogenesis, and demonstrates a graded pattern, this would suggest that this enhancer is required to promote or maintain *nr2f1a* expression within more venous ACs at later stages.

In summary, our data have provided insights into a Foxf-Wnt-Nr2f regulatory network that determines the number of differentiated ACs within the zebrafish heart. Our study has implications for our understanding of the development and evolution of gene regulatory networks controlling posterior SHF and AC differentiation in vertebrates and given the conserved requirements of Nr2f factors could be used to inform us of mechanisms underlying congenital heart defects that affect the atria in mammals. Future studies will be aimed at deciphering the specific transcriptional mechanisms that control the logic of this regulatory network and expanding our analysis to additional CREs directing *Nr2f1* and *Nr2f2* expression in ACs.

## Acknowledgements

We thank Kendall Martin for technical assistance, training, and reading of the manuscript.

## Materials and Methods

### Ethics statement

All zebrafish husbandry and experiments were performed following protocols approved by the Institutional Animal Care and Use Committee (IACUC) of Cincinnati Children’s Hospital Medical Center (Protocol IACUC2023-1048).

### Zebrafish husbandry and lines used

Adult zebrafish were raised and maintained under standard laboratory conditions [89]. WT fish used were mixed AB-TL background. Transgenic and mutant lines used were: *Tg(−1.5_nr2f1a:GFP)^ci1020^, Tg(−1.4_nr2f1a:GFP)^ci1021^, Tg(−0.7s_nr2f1a:GFP)^ci1022^, Tg(−0.7m_nr2f1a:GFP)^ci1023^, Tg(−0.7l_nr2f1a:GFP)^ci1024^, Tg(5’reg1-nr2f1a:GFP)^ci1025^, Tg(3’reg1-nr2f1a:GFP)^ci1026^, Tg(3’reg2-nr2f1a:GFP)^ci1027^, Tg(3’reg3-nr2f1a:GFP)^ci1028^, Tg(3’reg4-nr2f1a:GFP)^ci1029^, Tg(−5.1myl7:DsRed2-NLS)^f2^* [57], *Tg(−5.1myl7:EGFP)^twu26^* [90], *smo^s294^* [91], *nr2f1a^ci1009^* [35].

### Analysis of *Nr2f* loci and TFBS analysis

ATAC-seq data used in this study on flow-sorted zebrafish ACs at 48 hpf and alignments shown were reported previously (GEO, Accession # GSE 194054) [35]. Regions of open chromatin 700 kb upstream (a gene desert) and 18 kb downstream of the *nr2f1a* zebrafish locus were initially examined within the UCSC browser (https://genome.ucsc.edu). Open chromatin in these regions of the zebrafish genome adjacent to the *nr2f1a* locus that appeared to have conservation based on alignments with genomes available in the UCSC genome browser were selected for additional analysis of conservation through alignments using mVista (http://genome.lbl.gov/vista/mvista/submit.shtml) and subsequent reciprocal nBLASTs on Ensembl (Ensembl.org). Specific sequences from species corresponding to the conserved regions of open chromatin flanking *Nr2f1* loci in other species were retrieved from Ensembl: Australian ghostshark (*Callorhinchus*_*milii*-6.1.3), Tropical clawed frog (UCB_*Xtro*_10.0), Green anole (*AnoCar*2.0v2), Chicken (bGalGal1.mat.broiler.GRCg7b), House mouse (GRCm39), and Human (GRCh38.p14). The conserved regions in the different species, corresponding to the open chromatin in the zebrafish ACs, were manually aligned using Clustal Omega [92]. Sequences with a high degree of conservation from the alignment were analyzed for the presence of conserved TFBS using CIS-BP [40,41], TomTom [42,43], and JASPAR databases [44].

### Cloning of Plasmids

The *nr2f1a* promoter and enhancer reporters were generated using established Gateway cloning methods [93]. The designated regions of the *nr2f1a* promoter were cloned into *pDONR P4-P1r* to generate 5’-entry clones. Subsequently, each of the *p5E-nr2f1a* promoter clones were combined with *pME-EGFP* middle-entry and *p3E-polyA* 3’-entry clones into the previously reported *pDestTol2p2a-cry:DsRed* destination vector [93] to generate the *nr2f1a* promoter reporter constructs. The putative enhancer regions were cloned into the *pE1b-GFP-Tol2-Gateway* vector (Addgene, plasmid # 37846) [38]. A QuickChange Site-Directed Mutagenesis Kit (Agilent, Cat # 200524) was employed to generate the designated deletions and mutations of the putative binding sites within the *3’reg1-nr2f1a:GFP* reporter vector. Primers used for cloning are listed in **S2 Table**.

The murine *Foxf1* construct (gift of V. Kalinichenko) was reported previously [72]. The construct contains N-terminal Flag and C-terminal His tags. This construct was subcloned into *pCS2p+DEST1*, a version of *pCS2-Dest1* [94] with a Pst1 site added, using Gateway cloning. The *dnFoxf1* expression construct was generated via fusing the Engrailed transcriptional repressor domain with a 21 bp linker to the 5’-end of the murine *Foxf1* construct using PCR and placed into the *pCS2p+DEST1* vector.

### Morpholino (MO), mRNA, and plasmid injections

The *tcf7l1a* (aka *tcf3a*) MO was reported previously [95]. 1 ng of *tcf7l1a* MO was injected. mRNAs for *Foxf1*, *dnFoxf1*, and zebrafish *tcf7l1a* [54] were generated from linearized plasmid using a Sp6 Message Machine kit (Thermo-Fisher, Cat # AM1340). 50 pg of mRNA for *Foxf1* and *dnFoxf1* were injected. 5 pg of zebrafish *tcf7l1a* was injected. All injections were performed at the one-cell stage.

### Generation of transgenic lines

*Tol2* mRNA was generated from linearized plasmid using a Sp6 Message Machine kit (Thermo-Fisher, Cat # AM1340) [93]. For transient transgenesis and the generation of stable transgenic lines, 25 pg of plasmid DNA was co-injected with 25 pg of *Tol2* mRNA. For stable transgenic lines, fish were raised to adulthood and founders were identified via outcrossing to WT fish. The transgenic F1 fish were then selected and raised. At least two independent transgenic lines were initially examined and raised to confirm the equivalency of expression patterns. Only one transgenic line for each stable transgenic reporter was maintained. For experiments, homozygous transgenic *3’reg1:GFP* reporter fish were outcrossed to WT AB/TL fish.

### Drug treatments

All drug treatments were performed on embryos beginning at the 20s stage in 2 mL of embryo water with drugs at the specified concentrations in 3 mL glass vials. 30 embryos/vial were used for all experiments. Treated embryos in vials were placed on a nutator in an incubator at 28.5°C for the duration of the treatments. Concentrations of drugs used for treatments were: 1 μM RA (Sigma R2625), 2.5 μM DEAB (Sigma-Aldrich, Cat # D86256), 5 μM BIO (Tocris, Cat # 3194), 10 μM XAV939 (Tocris, Cat # 3748), 75 μM CYA (Sigma-Aldrich, Cat # C4116), 10 μM SAG (Tocris, Cat # 4366). All the drugs were rinsed out with blue water three times at 48 hpf prior to processing for IHC performed as described below.

### Immunohistochemistry (IHC) and cardiomyocyte quantification

IHC was performed as has been previously reported [18]. Briefly, embryos at the indicated stages were fixed for 1 hr at RT in 1% formaldehyde/1X PBS in 3 ml glass vials. Embryos were washed 1X in PBS and then 2X in 0.1% saponin/1X PBS, followed by blocking in 0.1% saponin/0.5% sheep serum/1X PBS (Saponin blocking solution) for one hr. Primary antibodies were applied to the embryos in blocking solution overnight at 4°C. Embryos were then gently washed 3X with 0.1% saponin/1X PBS followed by 2 hr incubation with secondary antibodies at RT. Secondary antibodies were washed out with 0.1% saponin/1X PBS. Primary antibodies used were: rabbit anti-Nr2f1a (1:100; custom antibody [12]), rabbit anti-DsRED (1:1000; Clontech, Cat # 632496), chicken anti-GFP (1:250; Invitrogen, Cat # A10262), mouse anti-Amhc (1:10, Developmental Studies Hybridoma Bank, Cat # S46), mouse anti-Mhc (1:10, Developmental Studies Hybridoma Bank, Cat #, MF20). Secondary antibodies used: Goat anti-mouse IgG1 DyLight (1:250, Biolegend, Cat # 409109), Goat anti-chicken IgY Alexa Fluor 488 (1:500, Invitrogen, Cat # A11039), Goat anti-rabbit IgG TRITC (1:100, Southern Biotech, Cat # 4050-03), Goat anti-mouse IgG2b TRITC (Southern Biotech, 1:100, Cat # 1090-03). Cardiomyocytes were quantified as previously reported [46]. Briefly, embryos were mounted ventral side down on coverslips. Images of the hearts were taken using a Nikon A1R Confocal Microscope with a 16X water immersion objective and the resonance scanner. The number of Nr2f1a^+^ and *myl7:*DsRed2-NLS^+^ nuclei in Amhc+ cardiomyocytes were counted in the images using Fiji [96]. Embryos were genotyped following imaging and cardiomyocyte quantification.

### In situ hybridization (ISH)

Whole-mount ISH was conducted similar to what has been previously reported [18]. Briefly, embryos were fixed overnight at 4°C in 4% paraformaldehyde/1X PBS in 3 ml glass vials. Subsequently, embryos were rinsed with PBS/0.1% Tween-20 (PBST) 3X and then dehydrated with increased percentage of methanol in PBST in series to 100% methanol and stored overnight at 4°C. Embryos were then rehydrated using a PBST/methanol series to 100% PBST, and then washed 4X in PBST. Digoxygenin-labeled *GFP* (ZDB-EFG-070117-2) riboprobe was used. Anti-Digoxygenin-AP, Fab fragment (Millipore-Sigma, Cat # 11093274910) was used to detect the Digoxygenin-labeled riboprobe in embryos. 3.5µL/mL NBT (Sigma-Roche, Cat # 11383213001) plus 3.5µL/mL BCIP (Sigma-Roche, Cat # 11383221001) in the Staining buffer (0.1M Tris-HCl, pH 9.5/0.1M MgCl2/1M NaCl/0.1% Tween-20) were used for the coloration reaction. Embryos were then rinsed with PBST 3X and dehydrated with a methanol/PBST series to 100% methanol, stored overnight at 4°C, followed by rehydration using a PBST/methanol series to 100% PBST. The embryos were then put in a glycerol/PBST series to 80% glycerol/PBST for imaging with a Zeiss_v12 M2Bio Stereo Microscope.

### Real Time qPCR

Total RNA isolation and RT-qPCR were performed as previously described [97]. Whole embryo RNA was obtained from groups of 30 24 hpf embryos using TRIzol (Invitrogen, Cat # 15596026) and Purelink RNA Microkit (Invitrogen, Cat # 12183016). cDNA was generated using 1μg total RNA and the Superscript Reverse Transcriptase kit (Invitrogen, Cat # 18080-051). RT-qPCR for *dkk1a* and *axin1*, was carried out using Power SYBR Green PCR Master Mix (Applied Biosystems, Cat # 4368706) in a BioRad CFX-96 PCR machine. Expression levels were standardized to *β-actin* expression [97] and data were analyzed using the Livak 2^−ΔΔCT^ Method [98]. Each experiment was performed in triplicate. Primers used are listed in **S3 Table.**

### Generation of zebrafish mutants and crispants

The *3’reg1* deletion mutants and *foxf1* crispants were made using CRISPR/Cas12a (Cfp1), similar to what has been reported [60]. *Acidaminococcus sp. BV3L6 (As)* crRNAs were generated to the 5’- and 3’-sequences of the *3’reg1-nr2f1a* enhancer and the coding region of *foxf1* exon1 using CHOPCHOP (http://chopchop.cbu.uib.no) [61]. The target-specific sequences at the 3’-end of the *As* crRNAs were fused with the common portion of the *As* crRNA with a T7 site at the 5’-end using PCR. The *As* crRNAs were then generated using T7 MEGAScript Kit (Thermo Fisher, Cat # AM1333). A solution of 2 *As* crRNAs (380 pg/µl) were incubated at RT with *As* Cas12a protein (2 µg/µl; IDT, Cat # 10001272) for 20 minutes prior to injection. 760 ng of the crRNA mixture was co-injected along with 4 ng of AsCas12a protein. Multiple pools of 10 embryos were screened for efficacy of the crRNA pairs in generating deletions with PCR. For the *3’reg1* enhancer, sibling embryos from injections that showed efficiency of the crRNAs in creating the appropriate deletions were raised to adulthood. Multiple pools of F1 progeny from founders were then screened for deletions using PCR, followed by cloning and sequencing. A *3’reg1* allele with a 354 bp deletion was identified and subsequently recovered. For the *foxf1* crispants, the *3’reg1:GFP* reporter embryos were scored for the presence of GFP and the individual embryos were screened for the efficacy of the deletions following imaging. Oligos used for generation of the sgRNA are listed in **S4 Table**. Primers used for genotyping and testing gRNA efficiency are listed in **S5 Table**.

### Statistical analysis

To determine if two proportions were statistically distinct, we performed Fisher’s exact test. To determine if proportions involving more than 2 conditions were statistically different, we performed a Chi-square test. To determine if the qPCR results were significant, we performed a Student’s t-test with Welch’s correction. To compare 3 or more distinct conditions (cell quantification), we performed an ordinary one-way ANOVA with multiple comparisons. Statistical analyses were performed utilizing GraphPad Prism. A p value <0.05 was considered statistically significant for all analysis.

