## Supporting Information - Figures and Figure Legends for "A Foxf1-Wnt-Nr2f1 cascade promotes atrial cardiomyocyte differentiation in zebrafish"

Supporting Information_Coppola et al.

### S1 Fig.

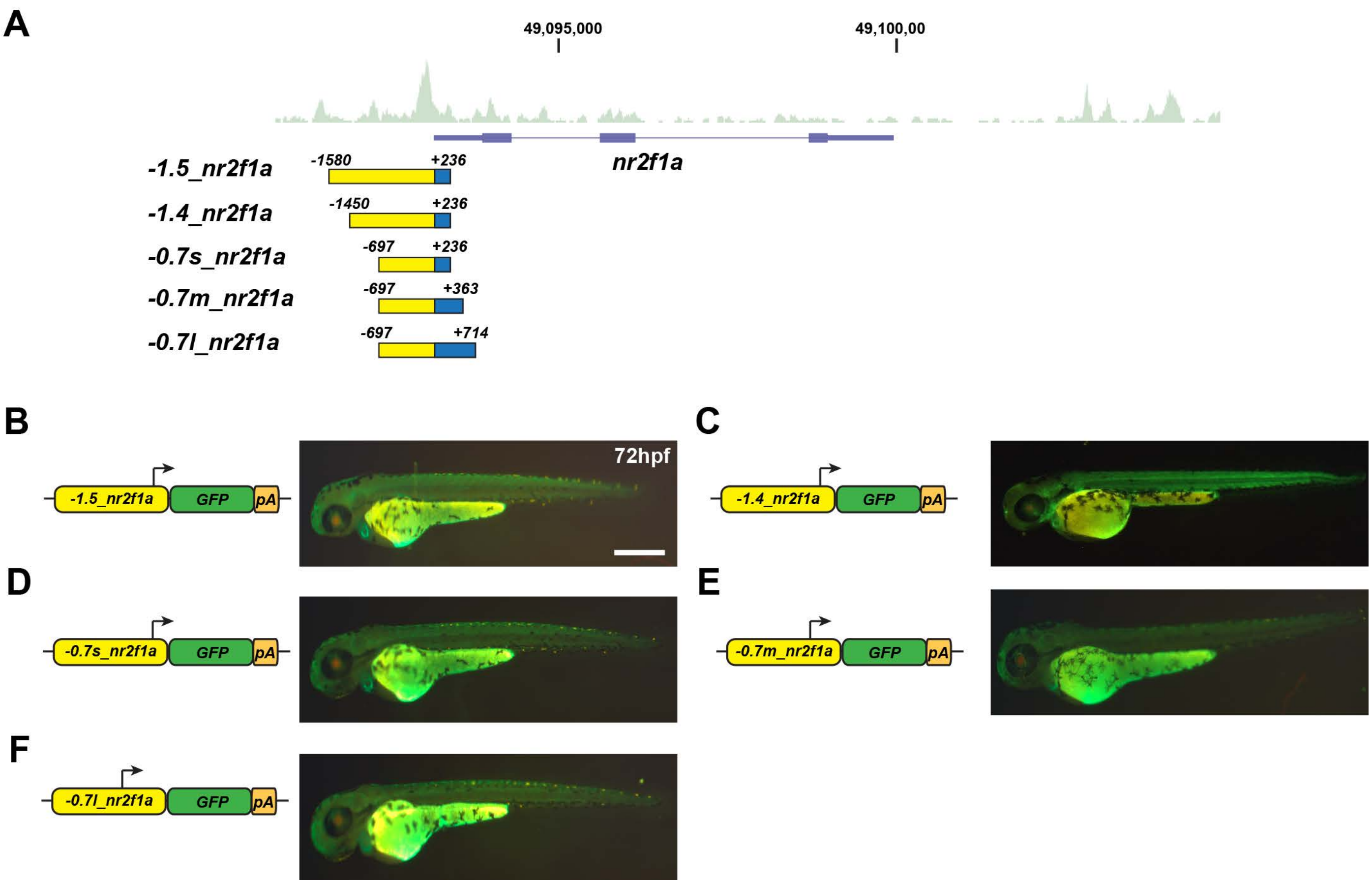


**S1 Fig. The proximal promoter region of *nr2f1a* promotes broad expression in zebrafish embryos. A**) Schematic of open chromatin in ACs from ATAC-seq showing the putative promoter fragments relative to the *nr2f1a* locus that were analyzed with stable transgenic lines. **B-F**) Schematics of the *nr2f1a* promoter transgenic constructs and images of representative stable transgenic lines. All the promoter constructs showed broad expression throughout the embryos. Lateral view with anterior left and dorsal upward. Scale bar: 500 μm.

**S2 Fig.**

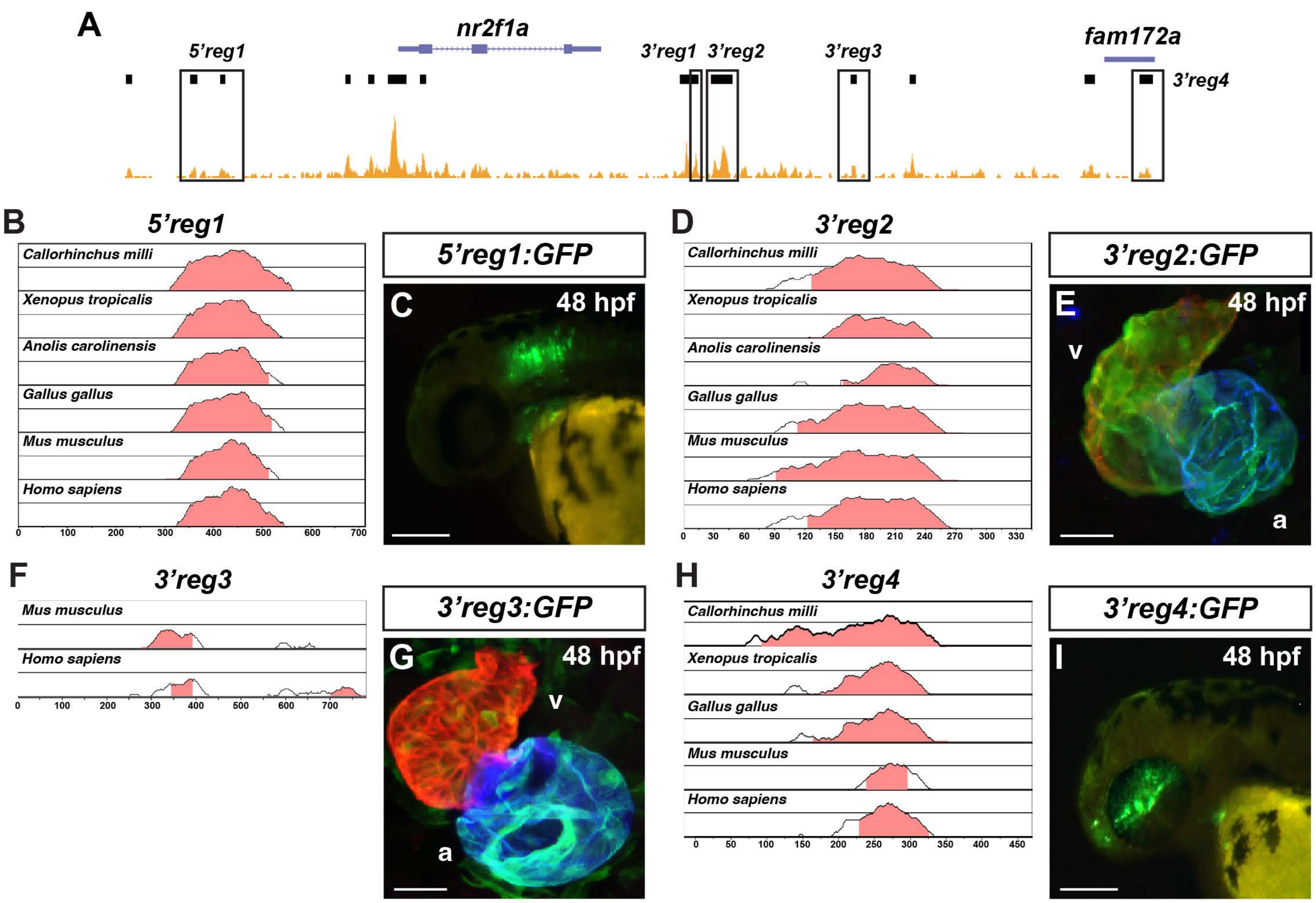


**S2 Fig. *Nr2f1a* enhancers promote expression in neural tissue and the heart. A**) Schematic of ATAC-seq data in ACs showing the localization of additional putative enhancers that were examined with reporters relative to the *nr2f1a* locus. **B**) VISTA plot showing conservation of the zebrafish *5'reg1-nr2f1a* enhancers with regions in *Callorhinchus milli* (Australian ghostshark), *Xenopus tropicalis* (Tropical clawed frog), *Anolis carolinensis* (Green Anole), *Gallus gallus* (chicken), *Mus musculus* (House mouse), and *Homo sapiens* (human). **C**) Image of *5'reg1:GFP* embryo with expression in the anterior hindbrain and branchial arches at 48 hpf. Lateral view is anterior to the left and dorsal upward. (n=25). Scale bar: 200 µm. **D**) VISTA plot showing conservation of the zebrafish *3'reg2-nr2f1a* enhancer with regions in *Callorhinchus milli* (Australian ghostshark), *Xenopus tropicalis* (Tropicalis clawed frog), *Anolis carolinensis* (green Anole), *Gallus gallus* (chicken), *Mus musculus* (House mouse), and *Homo sapiens* (human). **E**) Confocal image of heart at 48 hpf from *3'reg2:GFP* embryo stained for *3'reg2:GFP* (green), Vmhc (red), and Amhc (blue). V indicates ventricle. A indicates atrium. (n=21). Scale bar: 50 µm. **F**) VISTA plot showing conservation of the zebrafish *3'reg3-nr2f1a* enhancer with regions in *Mus musculus* (House mouse) and *Homo sapiens* (human). **G**) Confocal images of heart at 48 hpf from *3'reg3:GFP* embryos stained for *3'reg3:GFP* (green), Vmhc (red), and Amhc (blue). V indicates ventricle. A indicates atrium. Scale bar: 50 µm. **H**) VISTA plot showing conservation of the zebrafish *3'reg4-nr2f1a* enhancer with regions in *Callorhinchus milli* (Australian ghostshark), *Xenopus tropicalis* (Tropical clawed frog), *Gallus gallus* (chicken), *Mus musculus* (House mouse), and *Homo sapiens* (human). **I**) Image of the *3'reg4:GFP* embryo with expression in the medial eye, the anterior brain, the nasal pits, and the first branchial arch at 48 hpf (n=23). Scale bar: 200 µm. Pink in VISTA plots indicates >50% conservation of regulatory regions with the zebrafish enhancer sequence. Median lines in individual VISTA plots indicate 75% conservation. n indicates the number of embryos examined from a representative clutch. All images are of stable transgenic embryos.

**S3 Fig.**

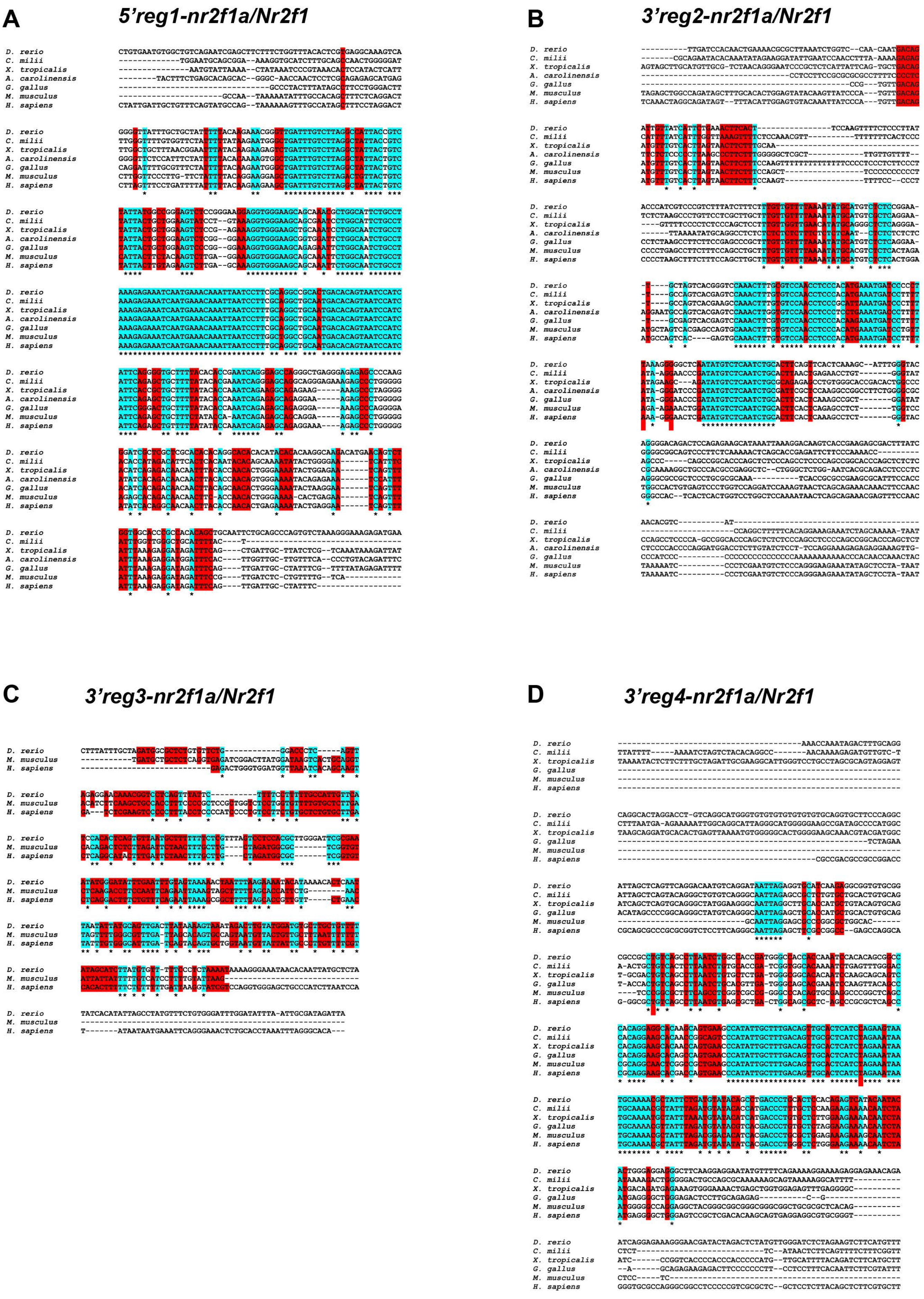
A
5'reg1-nr2f1a/Nr2f1
B
3'reg2-nr2f1a/Nr2f1
C
3'reg3-nr2f1a/Nr2f1
D
3'reg4-nr2f1a/Nr2f1
D. rerio
C. milii
X. tropicalis
A. carolinensis
G. gallus
M. musculus
H. sapiens

**S3 Fig. Conservation of *nr2f1a* enhancer sequences in other vertebrate species. A-D**) Clustal alignments of *5'reg1-nr2f1a/Nr2f1*, *3'reg2-nr2f1a/Nr2f1*, *3'reg3-nr2f1a/Nr2f1*, *3'reg4-nr2f1a/Nr2f1* between zebrafish and additional vertebrate species: *Callorhinchus milli* (Australian ghostshark), *Xenopus tropicalis* (Tropical clawed frog), *Gallus gallus* (chicken), *Anolis carolinensis* (Green Anole), *Mus musculus* (House mouse), and *Homo sapiens* (human). Turquoise indicates completely conserved nucleotides. Red indicates partially conserved nucleotides.

**S4 Fig.**

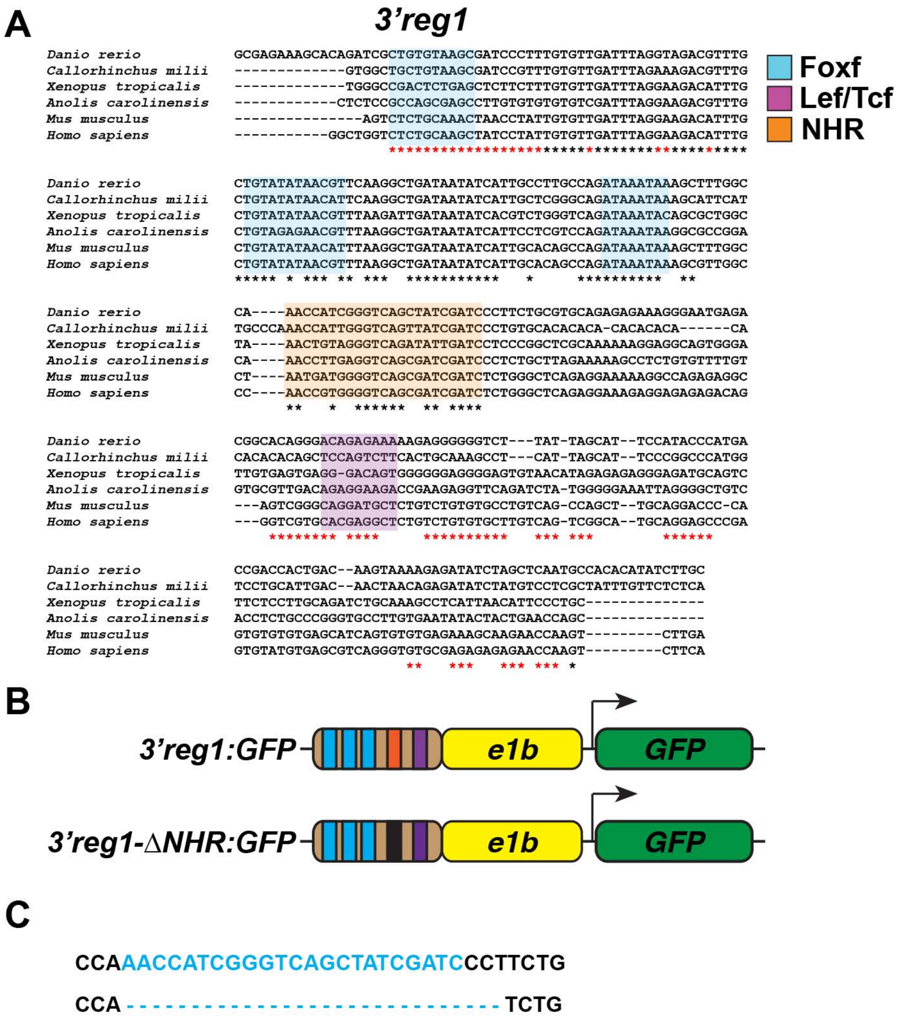


**S4 Fig. The NHR site within *3’reg1* is not required to promote expression or responsive to loss of RA signaling. A**) Clustal alignment of *3’reg1* with the putative NHR site highlighted, as well as Foxf1 and Lef/Tcf sites as shown in Fig. 1. **B**) Schematics of *3’reg1:GFP* reporter constructs. Foxf sites (blue), Lef/Tcf site (purple), NHR site (orange), deleted NHR site (black). **C**) Sequences showing WT NHR site and deletion of the NHR site in the *3’reg1:GFP* constructs. Deletion of the NHR site did not affect expression within the heart relative to the WT *3’reg1:GFP* construct (n=48) nor did treatment with the RA signaling inhibitor DEAB (N=33).

**S5 Fig.**

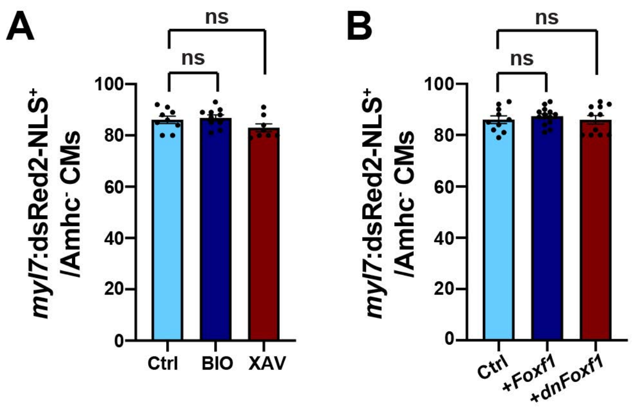


**S5 Fig. Wnt and Foxf1 manipulations do not impact VC number. A**) The number of VCs (*myl7*:DsRed2-NLS+/Amhc- cardiomyocytes) within the hearts of control, BIO-, and XAV-treated embryos. Control (n=9); BIO (n=10); XAV (n=8). **B**) The number of VCs (*myl7*:DsRed2-NLS+/Amhc- cardiomyocytes) within the hearts of control, *Foxf1* mRNA, and *dnFoxf1* mRNA-injected embryos. Control (n=10); *Foxf1* mRNA (n=13); *dnFoxf1* mRNA (n=11). Error bars in graphs indicate s.e.m..

**S6 Fig.**

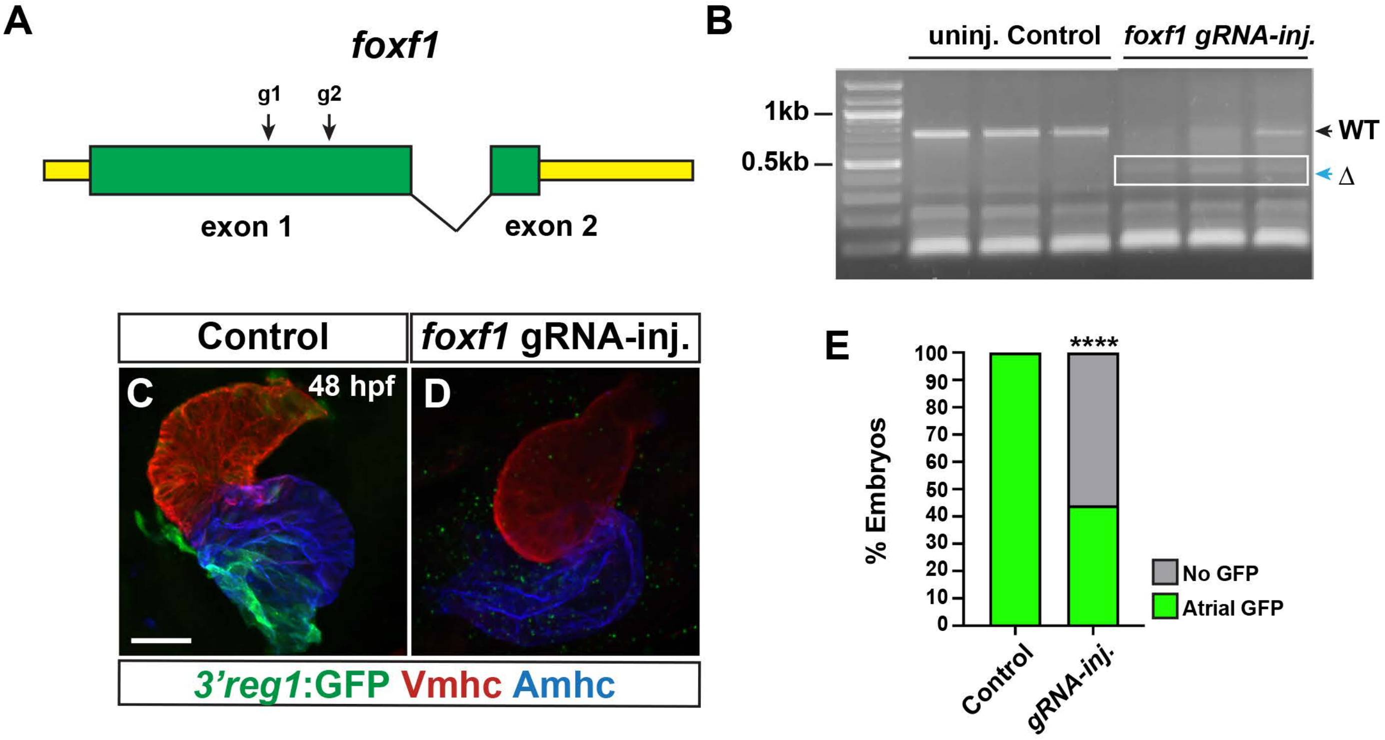


**S6 Fig. Loss of *foxf1* in zebrafish leads to a reduction in *3'reg1:GFP* expression within hearts.** **A**) Schematic showing the location of the guides (arrow) spaced ~200 bp apart in the first exon of zebrafish foxf1. **B**) PCR showing the efficacy of guides in creating an ~200 bp deletion and eliminating the WT band for foxf1 within representative injected *3'reg1:GFP* embryos. **C**) Confocal images of hearts from control and *foxf1* CRISPR-Cas12 injected transgenic *3'reg1:GFP* embryos stained for *3'reg1*:GFP (green), Vmhc (red), and Amhc (blue). Scale bars: 50 μm. D) The percentage of control uninjected and *foxf1* crispant *3'reg1:GFP* embryos with expression in the heart. Control (n=44); *foxf1* crispant (n=48). **** indicate $P < 0.0001$.

S7 Fig.

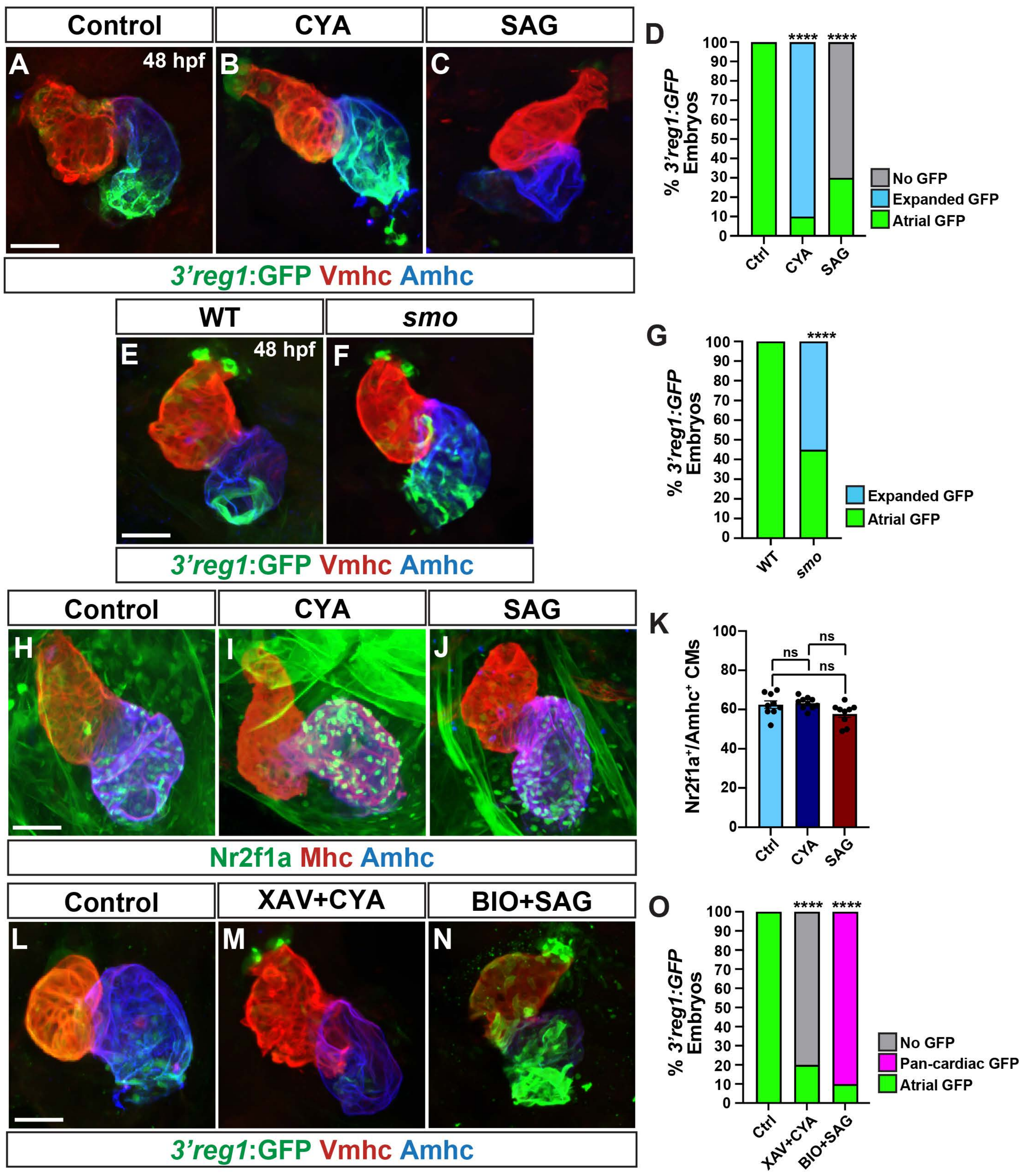

Control
CYA
SAG
A
48 hpf
B
C
D
% 3'reg1:GFP Embryos
No GFP
Expanded GFP
Atrial GFP
Ctrl
CYA
SAG
3'reg1:GFP Vmhc Amhc
WT
smo
E
48 hpf
F
G
Expanded GFP
Atrial GFP
Control
CYA
SAG
H
I
J
K
Nr2f1a+/Amhc+ CMs
ns
Nr2f1a Mhc Amhc
Control
XAV+CYA
BIO+SAG
L
M
N
O
No GFP
Pan-cardiac GFP
Atrial GFP
XAV+CYA
BIO+SAG

**S7 Fig. Hh signaling represses *3'reg1:GFP* expression within the hearts. A-C**) Confocal images of hearts from untreated control, CYA-treated, SAG-treated, XAV+CYA-treated, and BIO+SAG-treated transgenic *3'reg1:GFP* embryos stained for *3'reg1*:GFP (green), Vmhc (red), and Amhc (blue). **D**) The percentage of *3'reg1*:GFP embryos treated with CYA and SAG that had expanded or inhibited reporter expression within the atria. Control (n=31); CYA (n=44); SAG (n=98). **E,F**) Confocal images of hearts from WT sibling and *smo* mutant *3'reg1:GFP* embryos stained for *3'reg1*:GFP (green), Vmhc (red), and Amhc (blue). **G**) The percentage of WT and *smo 3'reg1*:GFP embryos that had expanded reporter expression within their atria. Control (n=24); *smo* (n=51). **H-J**) Confocal images of hearts from untreated control, CYA-treated, and SAG-treated stained for Nr2f1a (green), Mhc (red), and Amhc (blue). Scale bars: 50 µm. **K**) The number of Nr2f1a+/Amhc+ cardiomyocytes in the hearts of untreated control, CYA-treated, and SAG-treated embryos. Control (n=8); CYA (n=10); SAG (n=9). **L-N**) Confocal images of hearts from untreated control, XAV+CYA-treated, and BIO+SAG-treated transgenic *3'reg1:GFP* embryos stained for *3'reg1*:GFP (green), Vmhc (red), and Amhc (blue). **O**) The percentage of *3'reg1*:GFP embryos treated concurrently with XAV+CYA and BIO+SAG that had expanded or inhibited reporter expression within the atria. Control (n=23); CYA+XAV (n=99); SAG+BIO (n=109). Scale bars: 50 µm. Error bars in graph indicate s.e.m.. **** indicate $P < 0.0001$. ns indicates not a statistically significant difference.

**S8 Fig.**

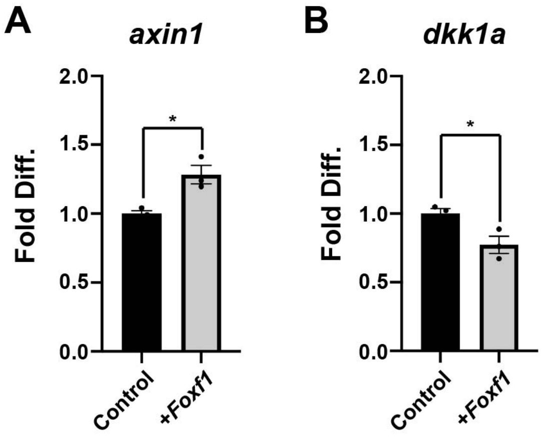


**S8 Fig. Foxf1 is sufficient to promote an increase in Wnt signaling in embryos at 24 hpf.** A,B) RT-qPCR for *axin1* and *dkk1a* in 24 hpf embryos injected with *Foxf1* mRNA. Fold difference is relative to *β-actin*. Error bars in graphs indicate s.e.m.. **** indicate P < 0.05.

**S9 Fig.**

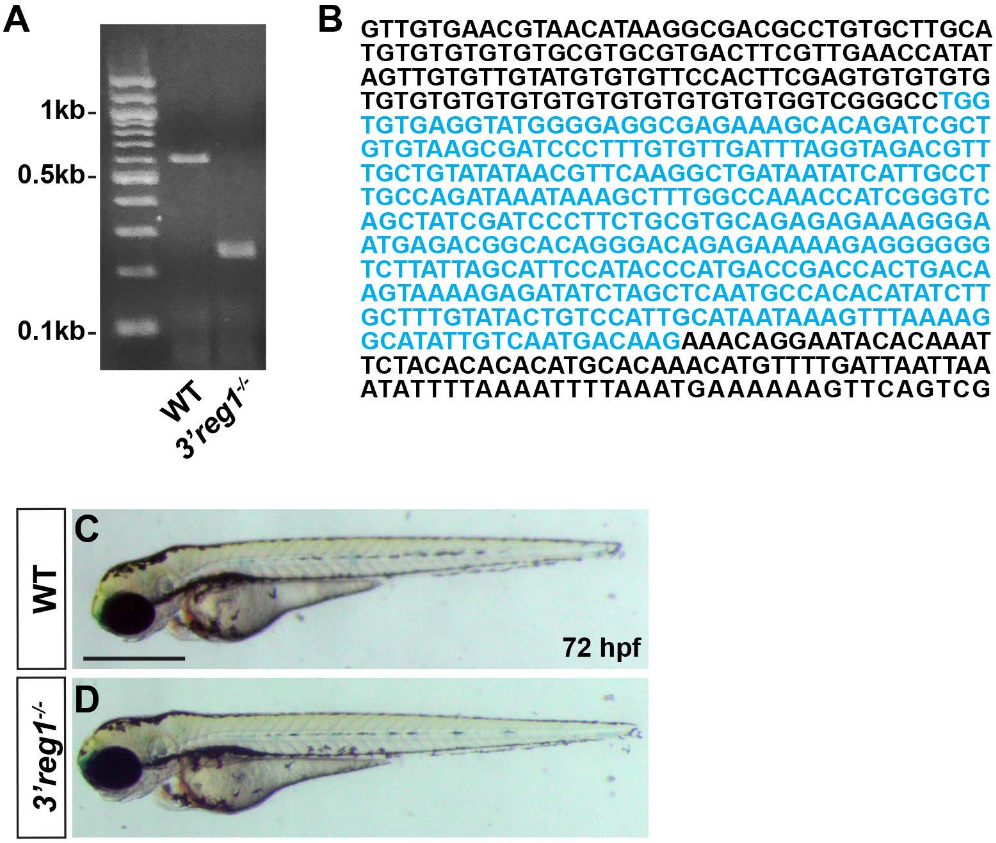


**S9 Fig. Zebrafish *3'reg1$^{-/-}$* embryos. A**) PCR from WT and *3'reg1* embryos. The WT *3'reg1* PCR product is 594 bp. The PCR product for the *3'reg1* deletion is 354 bp. **B**) *3'reg1* sequence showing the deleted sequence (blue). **C,D**) Representative WT and *3'reg1$^{-/-}$* embryos at 72 hpf. Lateral views with anterior leftward and dorsal upward. *3'reg1$^{-/-}$* embryos do not have an overt phenotype. Scale bar: 500 μm.

S10 Fig.

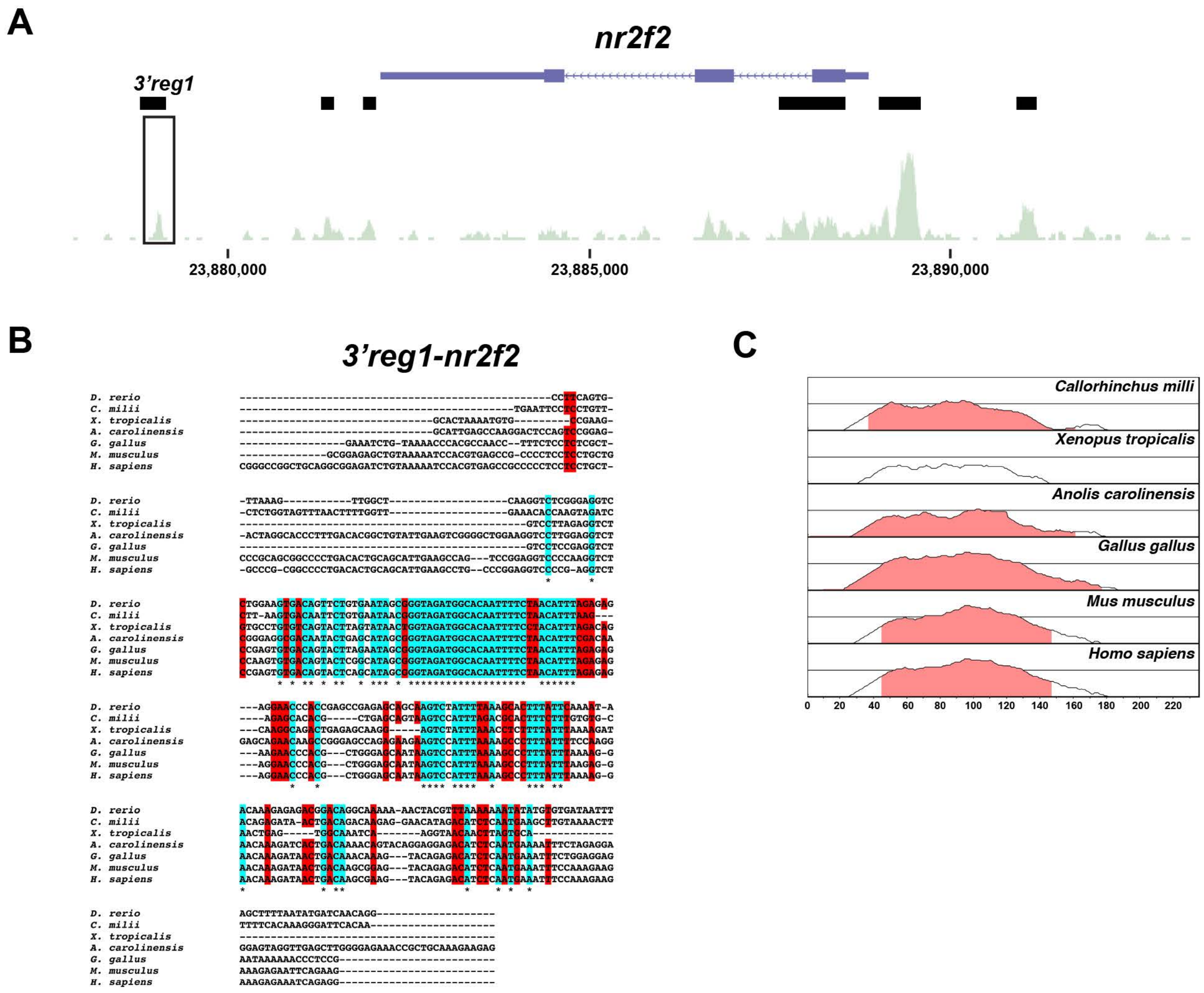


**S10 Fig. *3'reg1-nr2f2* enhancer is conserved in gnathostomes. A**) Schematic of ATAC-seq data in ACs showing the localization of the putative *3'reg1-nr2f2* enhancer relative to the *nr2f2* locus. **B**) Clustal alignment of *3'reg1-nr2f2* from *Danio rerio* (zebrafish), *Callorhinchus milli* (Australian ghostshark), *Xenopus tropicalis* (Tropical clawed frog), *Anolis carolinensis* (Green Anole), *Gallus gallus* (chicken), *Mus musculus* (House mouse), and *Homo sapiens* (human). Turquoise indicates completely conserved nucleotides. Red indicates partially conserved nucleotides. **C**) VISTA plot showing conservation of the zebrafish *3'reg1-nr2f2* enhancer with regions in *Callorhinchus milli* (Australian ghostshark), *Xenopus tropicalis* (Tropicalis clawed frog), *Anolis carolinensis* (green Anole), *Gallus gallus* (chicken), *Mus musculus* (House mouse), and *Homo sapiens* (human). Pink indicates >50% conservation of regulatory regions with zebrafish *3'reg1*. Median lines in individual VISTA plots indicate 75% conservation.
