## Supplementary material for "A Foxf1-Wnt-Nr2f1 cascade promotes atrial cardiomyocyte differentiation in zebrafish": S1 Table

**S1 Table. *Danio rerio* promoter sequences used in transgenic constructs. cloned in GFP vector with their names and genomic coordinates, respectively.**

>-1.5_nr2f1a_5:49743133-49744948

TTGTGTTCCCATTCGCTGTCCCTCTCCCTCTCTCCCCTCTCGCTCTCTTCCCTATCTGTGGAGTAAAACCTTGATGGTACACTTACAGAAGTTTCTCTTTGTTCGTTTTAATCTTATTCGTTTCACCTTTGCAATGAACAGAGGACAACAGCGTGTGTTTCTATGGATCTGTTTGTGGAGGGACGGAGAAGCTCCGCGGTCGAGTTGAAAATAAGCTACCTTTTTGTAACTAGAGGGTTTCTTGAACGGTCGGGTATCGGGGTTTTGTTTCCTCTTAACTTCACTTACACAGTTTTGGACTTTGATGCATTGTTTGTACAAAAAAAAAAAAAAACTTATGGTAAAAATAGGTGTTAGGTGATAAAGGCTACTAAAATATTTATTAAATACTTGAATTTTAAACATTGTTGTTTGTCCGCTAATAAGAGGCCAAAAAGCTTTTCTCCGAAAACAAAAATAAAATAAGCATCTCAAGTGGAGACATTGATAAAAAATAAAAATAAACTTGATGCTAAACAAGTTCCTGCAAAACATAATCAAAATACTTTTAAATCTGATGTTTTTAATAGCCTACTTCACTAAAAAAAATCAGAAACATTCATCCCACCGCGCAATTTGTCTACGAGTTACATTTGGAAACATGCTTTAAATGACAATAAATTACTAATTTTTCATGTCGCTGTAATTGTCTACTGTACGCTAGATGGCAGCCCGATCCAGACATTATAAGTCTGTACAGCGCAGCCTTCAGCATTCATGCTCAAAAAGCCGGTTTCCCTGCGCATTATACCAAAACTTCCCTCGTTAATAAATATGTTAAAACATGGATGGATCCATCGATGTATTTTTCTATTTTTCAATATCATTATACTTTGATTCCTCTTTTGTGCCCTGGTTTACTGCTTGATATGTTTCCTATCACTGAGAAAACATTCCGTTATAGCCTATACTGATATATCCCTCTAGTCAAACTAGTGGTAGTCATCGCGCTTCCCGGTGCTTTTTAATTTAATCGTTGTGCTGAGGATTAAGTGTAATATTGACACGTAAAAGCGAATTTTATACATCGATACGCAATTTCGAGTCGGTGGGCTATATCGGTCTGTCAAATTGCGTTAAACATTAATGGGACCTTCATAGTTCAAATCTGGACGCCGTTACGGTGTTTTTGTGTAATTAAGACTAGGGCTTTGAACAAAACACTTTCAACTTTGTGAGGTTCGCACCAGTCCTTTTGCCTCACACGCATTTGTTTATTTTTCACGTGCGCGTAGTTCGCTCATTGCTTTCATCCCGCTTTTCGCCGCTTCTTTCTGGGGTTTCTTCCTCTTTTTTTGTGACAATAGACGGTCCATTCTAATCGGGTTTCAAAGTAAATAGCAGCGCAGGGGCGAGAGCAATGTAAATTGTGCTTGTCAGAGCCCTCAGTCGTAGGTAGCGAGGAGCGAGGACTCTAACCAATGGAGTGAAGGAGGCTTGGCTAACCTTAGCCTCCCATTTTCTCTCTCCCCCCTCGATGCAGGGTTCTGACCAGTCAGTCGCCTTTGATTATCAGTTGCCAGCAGCCCCTCTCTTGGCTCCTTGACACGAGCACCATATAAGGCGCAGCGATCTCCATAGAAACGTGTCAGTTTCAATAGTAGTGTCAAAGTTCACTATATACAGACATTCGCGCAGATCTCCGTTTCGGAAACATTGCTCCGCTGGTACTCCCGTTAAAACGCATTCTTTTTGGGTCTCTGCTTCTTACATATTCCATTAGTTGCTTTTTTTCTTTTCTTTTCCTACTGGAGAGGTGAAACTACATAGCC

>-1.4_nr2f1a_5:49743263-49744948

GCAATGAACAGAGGACAACAGCGTGTGTTTCTATGGATCTGTTTGTGGAGGGACGGAGAAGCTCCGCGGTCGAGTTGAAAATAAGCTACCTTTTTGTAACTAGAGGGTTTCTTGAACGGTCGGGTATCGGGGTTTTGTTTCCTCTTAACTTCACTTACACAGTTTTGGACTTTGATGCATTGTTTGTACAAAAAAAAAAAAAACTTATGGTAAAAATAGGTGTTAGGTGATAAAGGCTACTAAAATATTTATTAAATACTTGAATTTTAAACATTGTTGTTTGTCCGCTAATAAGAGGCCAAAAAGCTTTTCTCCGAAAACAAAAATAAAATAAGCATCTCAAGTGGAGACATTGATAAAAAATAAAAATAAACTTGATGCTAAACAAGTTCCTGCAAAACATAATCAAAATACTTTTAAATCTGATGTTTTTATTAGCCTACTTCACTAAAAAAAATCAGAAACATTCATCCCACCGCGCAATTTGTCTACGAGTTACATTTGGAAACATGCTTTAAATGACAATAAATTACTAATTTTTCATGTCGCTGTAATTGTCTACTGTACGCTAGATGGCAGCCCGATCCAGACATTATAAGTCTGTACAGCGCAGCCTTCAGCATTCATGCTCAAAAAGCCGGTTTCCCTGCGCATTATACCAAAACTTCCCTCGTTAATAAATATGTTAAAACATGGATGGATCCATCGATGTATTTTTCTATTTTTCAATATCATTATACTTTGATTCCTCTTTTGTGCCCTGGTTTACTGCTTGATATGTTTCCTATCACTGAGAAAACATTCCGTTATAGCCTATACTGATATATCCCTCTAGTCAAACTAGTGGTAGTCATCGCGCTTCCCGGTGCTTTTTAATTTAATCGTTGTGCTGAGGATTAAGTGTAATATTGACACGTAAAAGCGAATTTTATACATCGATACGCAATTTCGAGTCGGTGGGCTATATCGGTCTGTCAAATTGCGTTAAACATTAATGGGACCTTCATAGTTCAAATCTGGACGCCGTTACGGTGTTTTTTTGTAATTAAGACTAGGGCTTTGAACAAAACACTTTCAACTTTGTGAGGTTCGCACCAGTCCTTTTGCCTCACACGCATTTGTTTATTTTTCACGTGCGCGTAGTTCGCTCATTGCTTTCATCCCGCTTTTCGCCGCTTCTTTCTGGGGTTTCTTCCTCTTTTTTTGTGACAATAGACGGTCCATTCTAATCGGGTTTCAAAGTAAATAGCAGCGCAGGGGCGAGAGCAATGTAAATTGTGCTTGTCAGAGCCCTCAGTCGTAGGTAGCGAGGAGCGAGGACTCTAACCAATGGAGTGAAGGAGGCTTGGCTAACCTTAGCCTCCCATTTTCTCTCTCCCCCCTCGATGCAGGGTTCTGACCAGTCAGTCGCCTTTGATTATCAGTTGCCAGCAGCCCCTCTCTTGGCTCCTTGACACGAGCACCATATAAGGCGCAGCGATCTCCATAGAAACGTGTCAGTTTCAATAGTAGTGTCAAAGTTCACTATATACAGACATTCGCGCAGATCTCCGTTTCGGAAACATTGCTCCGCTGGTACTCCCGTTAAAACGCATTCTTTTTGGGTCTCTGCTTCTTACATATTCCATTAGTTGCTTTTTTCTTTTCTTTTCCTACTGGAGAGGTGAAACTACATAGCC

>-0.7s_nr2f1a_5:49744016-49744948

CCTCTTTTGTGCCCTGGTTTACTGCTTGATATGTTTCCTATCACTGAGAAAACATTCCGTTATAGCCTATACTGATATATCCCTCTAGTCAAACTAGTGGTAGTCATCGCGCTTCCCGGTGCTTTTTAATTTAATCGTTGTGCTGAGGATTAAGTGTAATATTGACACGTAAAAGCGAATTTTATACATCGATACGCAATTTCGAGTCGGTGGGCTATATCGGTCTGTCAAATTGCGTTAAACATTAATGGGACCTTCATAGTTCAAATCTGGACGCCGTTACGGTGTTTTTGTGTAATTAAGACTAGGGCTTTGAACAAAACACTTTCAACTTTGTGAGGTTCGCACCAGTCCTTTTGCCTCACACGCATTTGTTTATTTTTCACGTGCGCGTAGTTCGCTCATTGCTTTCATCCCGCTTTTCGCCGCTTCTTTCTGGGGTTTCTTCCTCTTTTTTTGTGACAATAGACGGTCCATTCTAATCGGGTTTCAAAGTAAATAGCAGCGCAGGGGCGAGAGCAATGTAAATTGTGCTTGTCAGAGCCCTCAGTCGTAGGTAGCGAGGAGCGAGGACTCTAACCAATGGAGTGAAGGAGGCTTGGCTAACCTTAGCCTCCCATTTTCTCTCTCCCCCCTCGATGCAGGGTTCTGACCAGTCAGTCGCCTTTGATTATCAGTTGCCAGCAGCCCCTCTCTTGGCTCCTTGACACGAGCACCATATAAGGCGCAGCGATCTCCATAGAAACGTGTCAGTTTCAATAGTAGTGTCAAAGTTCACTATATACAGACATTCGCGCAGATCTCCGTTTCGGAAACATTGCTCCGCTGGTACTCCCGTTAAAACGCATTCTTTTTTGGGTCTCTGCTTCTTACATATTCCATTAGTTGCTTTTTTTCTTTTCTTTTCCTACTGGAGAGGTGAAACTACATAGCC

>-0.7m_nr2f1a_5:49744016-49745075

CCTCTTTTGTGCCCTGGTTTACTGCTTGATATGTTTCCTATCACTGAGAAAACATTCCGTTATAGCCTATACTGATATATCCCTCTAGTCAAACTAGTGGTAGTCATCGCGCTTCCCGGTGCTTTTTAATTTAATCGTTGTGCTGAGGATTAAGTGTAATATTGACACGTAAAAGCGAATTTTATACATCGATACGCAATTTCGAGTCGGTGGGCTATATCGGTCTGTCAAATTGCGTTAAACATTAATGGGACCTTCATAGTTCAAATCTGGACGCCGTTACGGTGTTTTTTTGTAATTAAGACTAGGGCTTTGAACAAAACACTTTCAACTTTGTGAGGTTCGCACCAGTCCTTTTGCCTCACACGCATTTGTTTATTTTTCACGTGCGCGTAGTTCGCTCATTGCTTTCATCCCGCTTTTCGCCGCTTCTTTCTGGGGTTTCTTCCTCTTTTTTTGTGACAATAGACGGTCCATTCTAATCGGGTTTCAAAGTAAATAGCAGCGCAGGGGCGAGAGCAATGTAAATTGTGCTTGTCAGAGCCCTCAGTCGTAGGTAGCGAGGAGCGAGGACTCTAACCAATGGAGTGAAGGAGGCTTGGCTAACCTTAGCCTCCCATTTTCTCTCTCCCCCCTCGATGCAGGGTTCTGACCAGTCAGTCGCCTTTGATTATCAGTTGCCAGCAGCCCCTCTCTTGGCTCCTTGACACGAGCACCATATAAGGCGCAGCGATCTCCATAGAAACGTGTCAGTTTCAATAGTAGTGTCAAAGTTCACTATATACAGACATTCGCGCAGATCTCCGTTTCGGAAACATTGCTCCGCTGGTACTCCCGTTAAAACGCATTCTTTTTTGGGTCTCTGCTTCTTACATATTCCATTAGTTGCTTTTTTTCTTTTCTTTTCCTACTGGAGAGGTGAAACTACATAGCCAGTCGGGCGACGTTGCTTTTTTTCGCTGGACCAGATGAGCTTTATTCATGAACATAGATAGAGAAATCCGTTCTTCAGTGTTTCCTCCTCTCCAACCGCGAAGACGGAGAGAGGAGCAAGGAAGAAA

>-0.7l_nr2f1a_5:49744016-49745426

CCTCTTTTGTGCCCTGGTTTACTGCTTGATATGTTTCCTATCACTGAGAAAACATTCCGTTATAGCCTATACTGATATATCCCTCTAGTCAAACTAGTGGTAGTCATCGCGCTTCCCGGTGCTTTTTAATTTAATCGTTGTGCTGAGGATTAAGTGTAATATTGACACGTAAAAGCGAATTTTATACATCGATACGCAATTTCGAGTCGGTGGGCTATATCGGTCTGTCAAATTGCGTTAAACATTAATGGGACCTTCATAGTTCAAATCTGGACGCCGTTACGGTGTTTTTGTGTAATTAAGACTAGGGCTTTGAACAAAACACTTTCAACTTTGTGAGGTTCGCACCAGTCCTTTTGCCTCACACGCATTTGTTTATTTTTCACGTGCGCGTAGTTCGCTCATTGCTTTCATCCCGCTTTTCGCCGCTTCTTTCTGGGGTTTCTTCCTCTTTTTTTGTGACAATAGACGGTCCATTCTAATCGGGTTTCAAAGTAAATAGCAGCGCAGGGGCGAGAGCAATGTAAATTGTGCTTGTCAGAGCCCTCAGTCGTAGGTAGCGAGGAGCGAGGACTCTAACCAATGGAGTGAAGGAGGCTTGGCTAACCTTAGCCTCCCATTTTCTCTCTCCCCCCTCGATGCAGGGTTCTGACCAGTCAGTCGCCTTTGATTATCAGTTGCCAGCAGCCCCTCTCTTGGCTCCTTGACACGAGCACCATATAAGGCGCAGCGATCTCCATAGAAACGTGTCAGTTTCAATAGTAGTGTCAAAGTTCACTATATACAGACATTCGCGCAGATCTCCGTTTCGGAAACATTGCTCCGCTGGTACTCCCGTTAAAACGCATTCTTTTTTGGGTCTCTGCTTCTTACATATTCCATTAGTTGCTTTTTTTCTTTTCTTTTCCTACTGGAGAGGTGAAACTACATAGCCAGTCGGGCGACGTTGCTTTTTTTCGCTGGACCAGATGAGCTTTATTCATGAACATAGATAGAGAAATCCGTTCTTCAGTGTTTCCTCCTCTCCAACCGCGAAGACGGAGAGAGGAGCAAGGAAGAAAAAAGAGGGGAATTTATTTTGCACAGCACTTTGGATCTGCGGTCCACCAGAAAGCTATTATTTTTGCTTCAACGTGAAGATTTTGTTTTTTACTGCGGTATTTTTTAAGAAAACTGTTTTTTTTATATTACGTCTGGGATCGCTTTCTTCATTCACGATTGGGTTCCCGAATGGCTGACTGCAATTTACCTTGGAACTGGCCTCCCGACAACTGCATATCCTGATCGGGTGCCTTTCTATCGACTCCGGTATTTTGAATGTATTGACCATTTTCTGCTTCTACTTTTTTCCCTATGAGATTGAGTGCTCCGATTTGAATTCGCGCTGCCGTTCGTCCAAGACTTCCCTTTT
